# When can a joint species distribution model capture biotic interactions? A simulation study across interaction mechanisms and levels of model integration

**DOI:** 10.64898/2026.05.20.726512

**Authors:** Jeanne Tous, Julien Chiquet

## Abstract

Joint species distribution models (JSDMs) describe the composition of communities by the response of each species to the environment and by the residual dependence between species, which can be read as a network of associations. These associations are not biotic interactions, but interactions are one of the processes that can produce them, and it remains unclear how much of a set of interactions an estimated network retains, and under which conditions. We address this question with a virtual ecologist approach. Communities are simulated along an environmental gradient with a known network of competition and facilitation, under three interaction mechanisms: a lottery for space (the VirtualCom model) and two models of population dynamics, Gompertz and Ricker. We compare four estimators of increasing integration, from a Gaussian graphical model on log-transformed counts to PLNnetwork, a Poisson log-normal model that estimates a sparse network of partial associations while it is fitted. Each estimated network is scored for two uses: ranking all the pairs of species, and shortlisting a few pairs. Each score is compared with the one obtained on a paired simulation without interactions, so that what an estimator owes to the niches alone is not counted. Within the assumptions of these simulators, five results emerge. (i) When the niches are narrow, interacting species seldom meet and there is little to capture, whatever the estimator. (ii) Partial associations do better than marginal ones, mostly among the best-ranked pairs. (iii) What a JSDM brings depends on the use and on the mechanism: its model for counts improves the ranking of all the pairs under the loglinear mechanism (Gompertz) only, mostly when counts are small or sampling effort is unequal, whereas estimating the sparse network together with the model improves the shortlist under the three mechanisms. (iv) Under the lottery and the Ricker mechanism, the Gaussian graphical model does as well over the whole ranking. (v) These gains require a covariate measured almost without error, no unmeasured gradient, a suitable representation of the abiotic effect of the covariate in the model, and enough latent factors. The scores remain modest throughout: a network inferred by a JSDM is at best a shortlist of candidate interactions, under conditions that this study makes explicit.

## 1. Introduction

In ecology, community data describe the distribution of a set of species in a set of sites, either as presence-absence, count or continuous data. These distributions result from three types of processes: dispersal, environmental filtering and biotic interactions (Guisan and Thuiller, 2005; D’Amen et al., 2018; Poggiato et al., 2021). Understanding these processes has been a long-term goal in ecology (Diamond, 1975; Connor and Simberloff, 1979) and can be critical to predict the responses of species communities to environmental stressors (Heinen et al., 2020).

However, these processes cannot be directly observed in general. To decipher species distribution data, a general methodology thus consists in identifying patterns in it (Gotelli and Graves, 1996; Ulrich and Gotelli, 2010), with the idea that these patterns can be linked to underlying processes or mechanisms. The complex relationship between processes, patterns and mechanisms has been the subject of a number of analyses (Anand, 1994; McAllister, 1997; Connolly et al., 2017). Anand (1994) proposes the following general definition: “patterns are what we perceive, processes describe how these patterns come about, and the mechanisms provide explanations as to why these patterns occur”. In this work, we focus on the relationship between patterns and processes, for one process, biotic interactions, and we vary the mechanism by which this process acts on the abundances of the species. Pattern detection is *per se* a difficult question, among other reasons because the definition of a pattern is both investigator and scale dependent (McAllister, 1997). The relationship between patterns and processes is also a complex one since different processes can have intermingled influences on species distributions and their effects may also be scale-dependent (Götzenberger et al., 2012; Viana and Chase, 2019). In our understanding, when a statistical model is fitted to community data, the patterns are the quantities that the model estimates, and the processes are what one hopes to read in them.

Historically, statistical ecology aimed at identifying and describing biodiversity patterns (Gimenez et al., 2014; Poggiato et al., 2021), using null models for instance (Connor and Simberloff, 1979; Ulrich and Gotelli, 2010). It has evolved towards more integrated modelling of ecological processes and the patterns they generate (Gimenez et al., 2014). Species Distribution Models (SDMs, Elith and Leathwick, 2009; Miller, 2010) and Joint Species Distribution Models (JSDMs, Pollock et al., 2014; Tikhonov et al., 2020; Chiquet et al., 2021) are two classes of models that belong to this integrated modelling of processes and patterns. SDMs model each species’ distribution data independently, relating it to abiotic covariates. Some of them additionally add other species as explanatory variable to model biotic interactions (Staniczenko et al., 2017; Poggiato et al., 2025) or integrate the modelling of dispersal effects (Shipley et al., 2022). JSDMs simultaneously model all the species in the data and consider both the effects of environmental covariates and the residual dependence between species, after accounting for the environmental covariates.

Most existing JSDMs, and all those discussed here, are formulated as hierarchical, or latent variable, models. Conditionally on a Gaussian latent vector that has one coordinate per species, the observations of the species at a site are independent, with a distribution suited to the data: Bernoulli for presence-absence data, which gives a probit model, or Poisson for counts. The latent vector carries what the covariates do not explain, and the dependence between species is described by its covariance matrix (Pollock et al., 2014; Ovaskainen and Abrego, 2020) or by the inverse of this matrix, the precision matrix (Chiquet et al., 2021). The model is the same in both cases; what differs is the parameter in which the network of species is read, and the two networks do not mean the same thing. In the first, two species are linked when they are correlated given the covariates, which corresponds to a *marginal association*. In the second, they are linked when they are still correlated given the covariates and all the other species, which corresponds to a *partial association*: a zero of the precision matrix means that the two species are independent given the others (Whittaker, 2009). Hence, two species that both respond to a third one have a marginal association and no partial one.

The initial interpretation given to these models was that, in both SDMs and JSDMs, the effects of covariates represented environmental filtering whereas, for JSDMs, residual correlations could be explained by biotic interactions, but also by missing predictors or model misspecifications (Warton et al., 2015). It is now clear that there is no equivalence between biotic interactions and associations in a JSDM-inferred network (Blanchet et al., 2020; Poggiato et al., 2021). This does not however deny that biotic interactions have a *possible role* as one of the factors that can explain the presence of a residual correlations between two species (Gotelli et al., 2010), and they are often invoked as one of the main ones (Pollock et al., 2014; Warton et al., 2015; Tikhonov et al., 2020). Still, it remains unclear which interactions leave a positive or a negative residual association, and under which conditions. We therefore study how much of a set of interactions the network estimated by a JSDM retains, depending on the interaction mechanism, the estimator, the data and the use made of the network.

Inferring associations between species from community data is older than JSDMs, and several families of methods address it. Graphical models estimate conditional dependences directly: Markov networks for presence-absence data (Harris, 2016), Bayesian networks (Milns et al., 2010), and Gaussian copula graphical models (Popovic et al., 2019), which accommodate counts as well as other types of data and are direct competitors of the Poisson log-normal models. Among JSDMs, the structure imposed on the dependence between species varies: latent factors, that is a covariance of low rank (Warton et al., 2015; Chiquet et al., 2018; Niku et al., 2019; Ovaskainen and Abrego, 2020); a covariance matrix estimated with a regularisation that induces sparsity, which gives a sparse network of marginal associations (Pichler and Hartig, 2021); a sparse precision matrix, which gives a sparse network of partial associations (Chiquet et al., 2019; Tous et al., 2026); and models built on deep neural networks, generic for multivariate binary responses (Chen et al., 2018) or designed for community data (Si-Moussi et al., 2026). Two results of this literature matter here: (i) partial associations separate direct effects from indirect ones (Harris, 2016; Popovic et al., 2019), and on simulated communities with known interactions the methods that control for indirect effects outperform those that do not (Harris, 2016); (ii) biotic interactions leave an imprint on co-occurrence and on spatial associations, which has been studied in theory (Cazelles et al., 2016), by comparing null-model, SDM-based and JSDM-based frameworks on empirical plant communities (D’Amen et al., 2018), and by comparing measures of association on empirical and simulated data (Keil et al., 2021). Yet the networks that the most widely used JSDMs report, such as gllvm (Niku et al., 2019) and Hmsc (Ovaskainen and Abrego, 2020), are networks of residual marginal correlations.

To explore this question, we propose to use a method in the spirit of the “virtual ecologist approach” (Zurell et al., 2010) that consists in testing methods (JSDMs here) on artificial data simulated from explicit rules of community assembly. The data are generated by simulators that are distinct from the statistical models fitted to them and in which the interactions are known.

Our study crosses four factors: (i) the *mechanism* by which the interactions act, either a lottery for space, as in the VirtualCom model (Münkemüller and Gallien, 2015), or one of two models of population dynamics, Gompertz and Ricker; (ii) the *room* that the niches leave to the interactions, set by the breadth of the niches and by how closely the interaction network follows niche proximity; (iii) the *estimator*, among four estimators of increasing *integration*, from a Gaussian graphical model on log-transformed counts to PLNnetwork (Chiquet et al., 2019), a Poisson log-normal model that estimates a sparse network of partial associations while it is fitted; and the *use* made of the estimated network, either ranking all the pairs of species from the most to the least likely to interact, or shortlisting a small number of pairs. The four estimators are ordered so that two successive ones differ by a single ingredient. All the scores are read against a paired simulation in which the interactions are switched off, so that what an estimator owes to the niches alone is not counted.

Our simulation study allows us to address five questions.

1. *How much is there to capture?* We expect little when the niches are narrow, because species that interact then seldom meet.
2. *Do partial associations capture more than marginal ones, on the same fit?* We expect so, since the simulated interactions are direct effects between two species, and since the literature reports it on other simulators (Harris, 2016).
3. *Does an integrated JSDM for counts capture more than a Gaussian graphical model on transformed counts?* We expect a competition to appear as a negative partial association and a facilitation as a positive one, and the integrated JSDM to do better when the simulator follows the model that the JSDM describes.
4. *What is an advantage due to: the model for counts, or the integration?* We expect the model for counts to matter where the counts are small.
5. *How do the sampling effort and the measurement of the environment affect these answers?* We expect an unequal sampling effort, an error on the covariate or an unmeasured gradient to leave structure in the residual associations that can be mistaken for interactions.

That partial associations separate direct from indirect effects is therefore not new. Our contribution is to determine under which mechanism, for which use, with which estimator and with which data an estimated network retains something of the interactions.

Below, we first present the three simulators, the four estimators, the scoring of an estimated network and the experimental design. We then report the results, following these questions. Finally, we discuss what they imply for the interpretation of the networks that JSDMs infer from real data.

## 2. Materials and methods

### 2.1. Simulating community data under three interaction mechanisms

We simulated community data with three *simulators* that use the same species, the same sites along one environmental gradient, the same Gaussian niches and the same interaction network. They differ in the *interaction mechanism*, that is the way interactions act on the counts, and, as a consequence, in the nature of the counts (Section 2.1.5). The three mechanisms are a lottery for space, as in the VirtualCom model (Münkemüller and Gallien, 2015); multispecies Gompertz dynamics (Ives et al., 2003); and Ricker dynamics (Ricker, 1954), a discrete-time Lotka–Volterra model. Throughout the paper, “simulator” refers to these data-generating processes, and “model” or “estimator” to the statistical methods applied.

VirtualCom directly encodes symmetric competition and facilitation between pairs of species, which makes the expected pattern explicit: two competitors should be found together less often than two species with the same niches and no interaction, and two facilitators more often. It is, however, one particular view of how interactions shape a community, in which the network only modulates the chance of occupying a vacated place. The two dynamic simulators add the other common view, in which interactions act on the growth of populations: a competitor lowers, and a facilitator raises, the *per-capita growth rate* of a species, that is the factor by which its abundance is multiplied from one time step to the next. Comparing the three shows whether a conclusion about an estimator belongs to the estimator or to the mechanism behind the data.

#### 2.1.1. Species, sites and interaction network

The environment was a single gradient on [0, 100], sampled at *n* = 50 sites regularly spaced along it; *x*_*i*_ denotes the position of site *i*. The units of the gradient are arbitrary: the niche breadth and the interaction threshold defined below are expressed in the same units, and what matters is their size relative to the length of the gradient and to the distance between two neighbouring sites, which is about 2. The species pool held *p* = 20 species. Each species *j* had a Gaussian niche described by two numbers, its optimum *θ*_*j*_ and a breadth *b* common to all species:

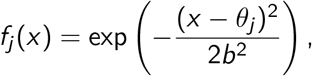

so that *f*_*j*_ equals 1 at the optimum. We drew the optima as VirtualCom does, through a simulated phylogeny: a tree is generated for the species of the pool by a pure-birth process, a trait evolves along its branches by Brownian motion, and the values of the trait at the tips, rescaled linearly to [0, 100], are the optima. Closely related species therefore have similar optima, and the optima are spread unevenly along the gradient, with groups of species sharing similar niches. Apart from this, the phylogeny plays no other role in the simulation. In particular, it enters neither the interaction network nor the estimators.

The interaction network is a symmetric matrix *W* = (*W*_*jk*_) with a zero diagonal, such that *W*_*jk*_ > 0 for two species in competition, *W*_*jk*_ < 0 for two species in facilitation, and *W*_*jk*_ = 0 otherwise. In the original model, any two species have a competition interaction, whose strength is directly related to the proximity of their niches. The interaction network is then full, of a single sign, and entirely determined by the niches. We chose instead a sparse network with interactions of both signs, drawn partly at random, for two reasons: first, we regard it as closer to real communities, where a species interacts with a few of the others and not with every species that shares its habitat; and second, an estimator can be scored on whether it tells the interacting pairs from the others only if some pairs do not interact, and if the network is not a function of the niches, which the covariate already describes. Therefore, we first fixed a distance threshold between niche optima, beyond which two species have no interaction: only the pairs with |*θ*_*j*_ − *θ*_*k*_| < *D* are eligible, *D* being the *interaction threshold*. Among the eligible pairs, we drew at random a fraction *d*_comp_ = 0.2 to be in competition, with *W*_*jk*_ = *a*, and a disjoint fraction *d*_faci_ = 0.1 to be in facilitation, with *W*_*jk*_ = −*a*. Every network therefore mixes the two types of interaction, and all the interactions of a network have the same intensity *a*.

Both the niche breadth and the interaction threshold decide how much room the niches leave to the interactions. The niche breadth *b* ∈ {2, 5, 10, 20} sets how many species coexist at a site, and hence how often interacting species meet; the interaction threshold *D* ∈ {10, 50} sets how far apart the optima of two interacting species can be: at *D* = 10 interacting species have nearly the same optimum, while at *D* = 50 the network is largely unrelated to the distance between optima. Whether two interacting species meet then depends on this distance relative to the niche breadth, that is on *D* and *b* together. We consider the eight *configurations* obtained by crossing the values of *b* and *D*. A network had on average 7 competition and 3 facilitation interactions among the 190 pairs of species at *D* = 10, and 29 and 14 at *D* = 50.

#### 2.1.2. Lottery for space (VirtualCom)

VirtualCom stochastically simulates community assembly at each site separately, as a lottery for the *K* places of the site, *K* being a carrying capacity that is the same for all sites (Supplementary material, Figure S1). We used this lottery as defined by Münkemüller and Gallien (2015), in their reference implementation, to which we only added the interaction network. At each site, the simulation proceeds as follows:

1. *Initialisation*. The *K* places of the site are filled with individuals whose species are drawn uniformly at random from the species pool.
2. *Replacement*. One individual of the site, chosen uniformly at random, is removed, and its place is taken by a new individual. The species of the newcomer is drawn with a probability that depends on the niche of the species, on its interactions with the species present at the site, and on its own local abundance (the expression of this probability is given below). The newcomer can be of the same species as the individual it replaces.
3. *Iteration*. Step 2 is repeated *K* times, which makes one “year” in the terminology of the model, and the simulation is run for several years, so that the community no longer depends on its random initial composition.

At step 2, the probability that the vacated place at site *i* is taken by an individual of species *j* is

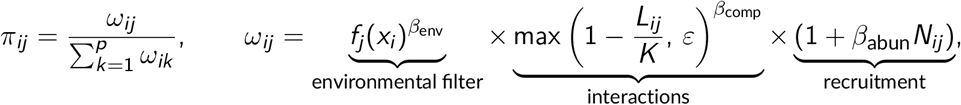

where *N*_*ij*_ is the current number of individuals of species *j* at site *i* and *L*_*ij*_ = ∑_*k*_ *W*_*kj*_ *N*_*ik*_ is the interaction load that the individuals present at the site exert on species *j*. The weight *ω*_*ij*_ is the product of the three factors of VirtualCom: the environmental filter, the interactions, and a recruitment that increases with the local abundance of the species. The competitors of *j* raise the load *L*_*ij*_ and lower the chance that *j* takes the place, and its facilitators do the opposite: with *a* = 2, a competitor holding half of the places excludes species *j*, and a facilitator of the same abundance doubles its weight. The constant *ε* = 10^−5^ is a safeguard that we added to keep the weight positive when the load exceeds *K*. The exponents *β*_env_, *β*_comp_ and *β*_abun_ weight the three factors; all were set to 1. We used *K* = 100 individuals per site, 10 years (Supplementary Section S4, Figure S4), and three interaction intensities, *a* ∈ {0.5, 2, 5}.

We wish to underline two properties of this mechanism that matter for what follows. First, the network acts only on the probability of winning a place, hence the lottery reproduces the consequence that an interaction is expected to have on the composition of a community, and it does not describe a process acting on the growth of populations. Second, the community is closed. Indeed, each site holds exactly *K* individuals, so that all the species compete for space whatever the network, and the counts are an exact census of the site.

#### 2.1.3. Gompertz dynamics

Under the Gompertz mechanism, the log-abundance of each species at a site fluctuates around a level set by the environment, *µ*_*ij*_ = log *K* + *β*_env_ log *f*_*j*_ (*x*_*i*_). The vector *d*_*i*_ (*t*) ∈ ℝ ^*p*^ of the deviations from this level follows a first-order multivariate autoregressive process (Ives et al., 2003), independently at each site:

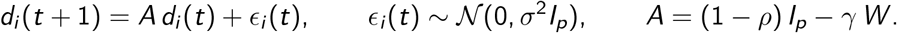

Here *ρ* is the rate at which a species returns to its environmental level (its self-regulation), *σ* is the process noise and *γ* scales the interactions. A competitor of species *j* that stands above its own level lowers the next log-abundance of *j*, and a facilitator raises it. Starting from *d*_*i*_ (0) = 0, we ran *T* = 200 steps and drew the counts as *Y*_*ij*_ ~ *P* (exp(*µ*_*ij*_ + *d*_*ij*_ (*T*))).

For every network to give a stationary process, we set the strength of the interactions relative to self-regulation, *γ* = *β*_comp_ *κρ/λ*_max_(*W*), where *λ*_max_(*W*) is the largest absolute eigenvalue of *W* and *κ* ∈ (0, 1) is the *interaction strength*. Since *W* = *aS*, where *S* holds the signs of the interactions, *λ*_max_(*W*) = *aλ*_max_(*S*): the intensity *a* cancels out, and only *κ* matters. We used *ρ* = 0.5, *κ* ∈ {0.5, 0.9} and *σ* ∈ {0.3, 0.5, 0.8}.

This mechanism is log-linear, which gives it a special place. For a symmetric *W*, the stationary covariance of the deviations solves Σ = *A*Σ*A*^*⊤*^ + *σ*^2^*I*_*p*_ (Ives et al., 2003), that is Σ = *σ*^2^(*I*_*p*_ −*A*^2^)^−1^, and a direct calculation gives their precision matrix:

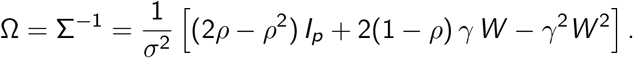

The data therefore follow a Poisson log-normal distribution, in which the environmental level is quadratic in *x*_*i*_ and the interaction network is written, to first order in *γ*, in the precision matrix of the Gaussian layer: a competition gives a positive entry of Ω, that is a negative partial correlation, and a facilitation a negative entry. The term in *W* ^2^, of second order in *γ*, adds entries between two species that share a partner. These are not interactions of *W*, and they count as errors when an estimated network is compared with *W*.

#### 2.1.4. Ricker dynamics

The Ricker mechanism is the multispecies form of the model of Ricker (1954). The environment enters through a carrying capacity shaped by the niche, 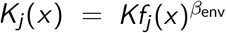, around which the density of species *j* at site *i* fluctuates in the absence of interactions; the interactions act on per-capita growth relative to this capacity, on the scale of abundances:

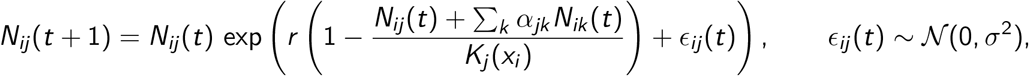

with *α* = *β*_comp_ *κ W /λ*_max_(*W*). Here *N*_*ij*_ (*t*) is a continuous density, the number of individuals expected in the sampling unit, and not a number of individuals. Starting from *N*_*ij*_ (0) = *K*_*j*_ (*x*_*i*_)*/*2, we ran *T* = 200 steps with an intrinsic growth rate *r* = 0.5 and drew the counts as *Y*_*ij*_ ~ *P*(*N*_*ij*_ (*T*)): the expected count follows the niche through *K*_*j*_ (*x*_*i*_). The interaction strength *κ* and the process noise *σ* took the same values as under the Gompertz mechanism.

This mechanism is not log-linear: competitive exclusion and absences at the edges of the niches come out of the dynamics. Apart from log-linearity, it shares everything with the Gompertz mechanism.

#### 2.1.5. Comparison of the three mechanisms

Table 1 sets the three mechanisms side by side. Two differences separate the lottery from the Gompertz mechanism: the form of the interactions and the nature of the data (an exact census of a closed community for the former and a Poisson sample of an open community for the latter). The Ricker mechanism makes the link between the lottery and the Gompertz mechanism: it has the per-capita effects, the strength, the noise and the Poisson observation of the Gompertz mechanism, and differs from it only by log-linearity.

**Table 1.** The three interaction mechanisms. A setting is a configuration (niche breadth and interaction threshold), a mechanism and a strength of the interactions.

|  | Lottery | Gompertz | Ricker |
|---|---|---|---|
| Interactions act on | the probability of taking a vacated place | per-capita growth, on the log scale | per-capita growth, on the scale of abundances |
| Log-linear | no | yes | no |
| Network in the precision matrix of a latent Gaussian layer | no | yes, to first order | no |
| Individuals per site | fixed (*K*) | free | free |
| Counts | exact census | Poisson sample of a density | Poisson sample of a density |
| Strength of the interactions | intensity *a* ∈ {0.5, 2, 5} | *κ* ∈ {0.5, 0.9}, with *σ* ∈ {0.3, 0.5, 0.8} | *κ* ∈ {0.5, 0.9}, with *σ* ∈ {0.3, 0.5, 0.8} |
| Settings | 8 × 3 = 24 | 8 × 6 = 48 | 8 × 6 = 48 |

All three simulators rest on the same simplifying assumptions: one environmental gradient, observed without error; Gaussian niches of equal breadth; symmetric interactions between pairs of species, all of the same intensity; and no dispersal between sites. Some of them are relaxed in separate experiments (Section 2.4).

#### 2.1.6. Replicates and paired no-interaction control

We consider 120 *settings*, obtained by crossing the configurations, the mechanisms and the strengths of the interactions (Table 1). Each setting was replicated 40 times, each replicate drawing its own species pool and its own network. Replicate number *r* of a configuration has the same niche optima and the same network under the lottery at *a* = 2 and under the Gompertz and Ricker mechanisms at every *κ* and *σ*, so that the mechanisms are compared on the same communities.

Every replicate was simulated a second time from the same random state with the interactions switched off (*β*_comp_ = 0): this *paired no-interaction control* differs from the simulation with interactions only in that the network does not act.

A simulation outputs the counts *Y*_*ij*_ of the *p* species at the *n* sites, the niche optima and the interaction network. Species absent from all the sites were removed, and the control was restricted to the species kept in the simulation with interactions. The simulated data are described in Supplementary Section S2.

#### 2.1.7. Unequal sampling effort

In a separate experiment, the sites were sampled with unequal effort. Each site *i* received an effort *e*_*i*_ ∈ (0, 1], the probability that an individual present at the site is counted, with 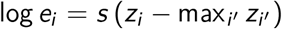 and *z*_*i*_ ~ *N* (0, 1). The observed counts were obtained by binomial thinning of the simulated ones, 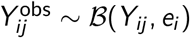. The *spread s* sets how unequal the effort is. A simulation and its control share the same efforts, as do all the values of *s*.

### 2.2. Estimating the network: four levels of integration

All the estimators considered separate what the environment explains in the counts from the dependence between species, and read a sparse network in the latter. They differ in how each step is carried out and in how tightly the two are tied together. We arranged them in four *levels of integration*, such that two successive levels differ by one ingredient only (Table 2). Their mathematical form and their settings are given in Supplementary Section S3.

**Table 2.** The four levels of integration. From level 1 to level 3 the estimator gains a model for counts; from level 2 to level 3 it loses a constraint on the rank; from level 3 to level 4 the sparse network is estimated together with the rest of the model.

| Level | Estimator | Model for the counts | Latent covariance | Sparse network |
|---|---|---|---|---|
| 1 | log-GGM | Gaussian, on log(1 + *Y*) | full rank | graphical lasso on the estimated covariance (two-step) |
| 2 | PLNPCA + graphical lasso | Poisson log-normal | rank 10 | graphical lasso on the estimated covariance (two-step) |
| 3 | PLN + graphical lasso | Poisson log-normal | full rank | graphical lasso on the estimated covariance (two-step) |
| 4 | PLNnetwork | Poisson log-normal | sparse precision | estimated with the model (integrated) |

#### 2.2.1. The common framework

The four levels are versions, exact or approximate, of one joint species distribution model, the Poisson log-normal model (PLN, Aitchison and Ho, 1989; Chiquet et al., 2021). Given a Gaussian latent vector, the counts of the species at a site are independent Poisson variables:

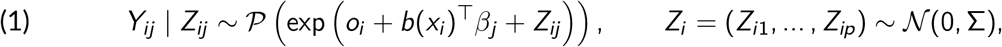

independently from one site to the next. Here *b*(*x*_*i*_) is the vector of the *abiotic basis* at site *i*, a set of functions of the gradient described below; *β*_*j*_ holds the coefficients of species *j*, which are specific to each species and share nothing between species; and *o*_*i*_ is an offset, equal to zero unless stated. The term *b*(*x*_*i*_)^⊤^*β*_*j*_ is the abiotic effect. The latent vector *Z*_*i*_ carries what the environment does not explain, and its covariance matrix Σ, or the precision matrix Ω = Σ^−1^, describes how the species vary together once the abiotic effect is removed.

As in the introduction, the *residual associations* are *marginal* when read in Σ and *partial* when read in Ω. Since the partial correlation between *Z*_*ij*_ and *Z*_*ik*_ is 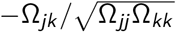 (Whittaker, 2009), a pair of species in competition is expected to give Ω_*jk*_ > 0 and a pair in facilitation Ω_*jk*_ < 0, the pattern that the three simulators are meant to produce (Section 2.1).

#### 2.2.2. The four levels of integration

The four levels are the following.

1. *The log-GGM* is a Gaussian graphical model on covariate-adjusted log counts. The counts are transformed to log(1+*Y*_*ij*_) and regressed on the abiotic basis by least squares, species by species; the covariance of the residuals is computed; and the graphical lasso (Friedman et al., 2008) turns it into a sparse precision matrix. This is model (1) in which the Poisson layer is replaced by a transformation of the data, that is, an approximate JSDM by Gaussian graphical modelling that makes a direct competitor to PLNnetwork.
2. *PLNPCA followed by the graphical lasso* fits model (1) with a covariance of reduced rank, Σ = *CC* ^⊤^ with *C* of size *p*×*q* (Chiquet et al., 2018). This is the latent-factor JSDM: the *p* species load on *q* latent factors, as in gllvm and Hmsc. The graphical lasso is then applied to the rank-*q* estimate of Σ returned by PLNPCA returns (Supplementary Section S3). We used *q* = 10 in the comparison of the levels, and fitted *q* = 2 and *q* = 5 as well to study the effect of the rank, 2 being the default number of latent variables of gllvm.
3. *PLN followed by the graphical lasso* fits model (1) with an unconstrained Σ, and applies the graphical lasso to its estimate.
4. *PLNnetwork* estimates a sparse Ω together with the abiotic coefficients, in one step, by maximising a variational lower bound of the likelihood of model (1) penalised by the *l*_1_ norm of the off-diagonal entries of Ω (Chiquet et al., 2019).

Levels 1 to 3 are *two-step* estimators, a covariance first and a sparse precision matrix next; level 4 is *integrated*. Each step between two levels isolates one ingredient, with the same abiotic basis and the same graphical lasso on both sides. From level 1 to level 3, the transformation gives way to a model for counts. From level 2 to level 3, the constraint on the rank is lifted. From level 3 to level 4, the sparse network is no longer plugged in after the fit but estimated with it.

We chose the PLN framework to build the four levels because its variants share the same model, the same variational inference and the same implementation, in the R package PLNmodels: a difference between two levels can then be attributed to a modelling choice, and not to an inference method or to a piece of software. We do not claim that it is the only possible framework. The two most widely used JSDMs, gllvm (Niku et al., 2019; Korhonen et al., 2025) and Hmsc (Ovaskainen and Abrego, 2020), in their Poisson versions, are latent-factor models: they share the structure of level 2 and differ from it by the inference method. They are considered in Supplementary Section S6.

We complete the comparison with two controls. First, for levels 1 and 3 we also read the network in the covariance itself, by ranking the pairs on their residual correlation: partial and marginal associations are then compared on the same fit. Second, gllvm was fitted with a Poisson distribution, its default of two latent variables and its default variational approximation, and its residual covariance was put through the same graphical lasso as levels 1 to 3, so that it can be read in partial associations and compared with level 2 at the same rank.

#### 2.2.3. Five abiotic bases

The gradient enters the model via the abiotic basis *b*(*x*). This modelling choice is offered in the same terms to every estimator. The five variants compared are given in Table 3.

**Table 3.**
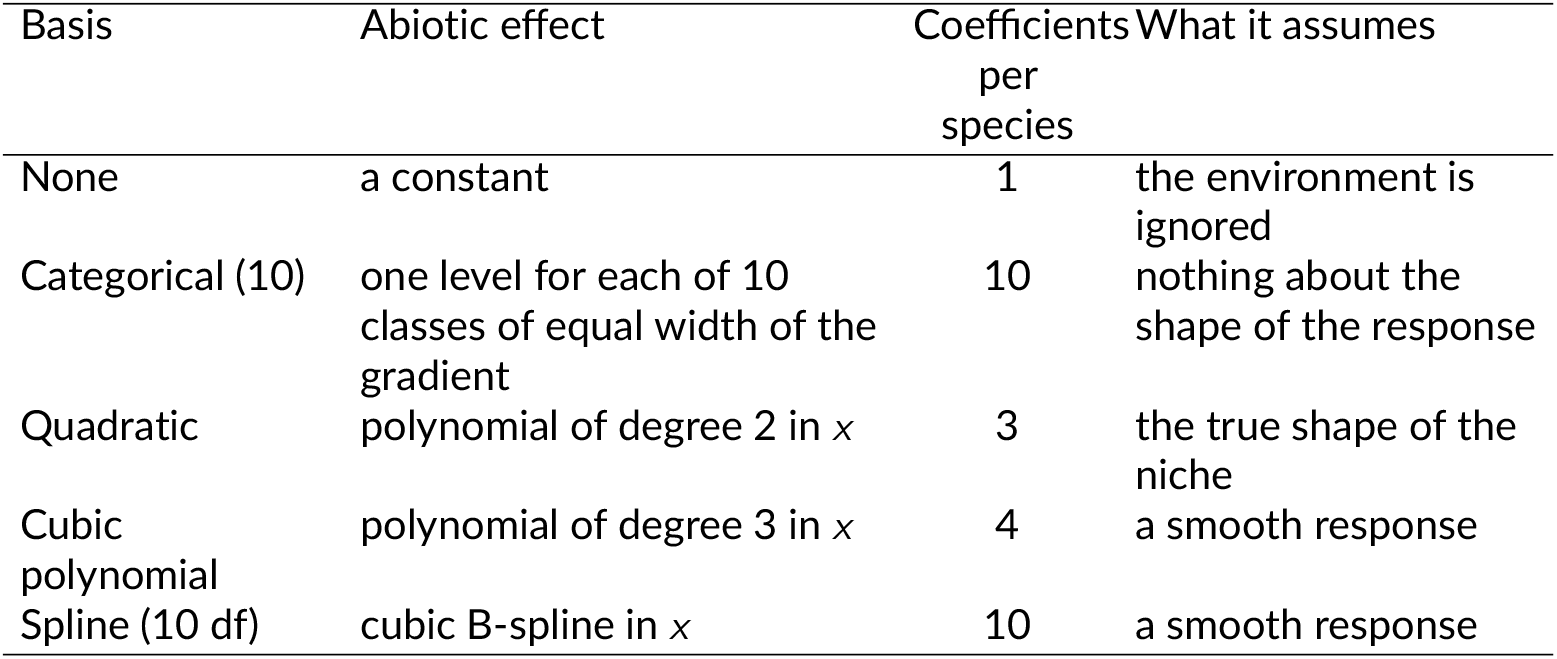
The five abiotic bases. The coefficients are counted with the intercept. The last three are the *smooth bases*.

| Basis | Abiotic effect | Coefficients per species | What it assumes |
|---|---|---|---|
| None | a constant | 1 | the environment is ignored |
| Categorical (10) | one level for each of 10 classes of equal width of the gradient | 10 | nothing about the shape of the response |
| Quadratic | polynomial of degree 2 in *x* | 3 | the true shape of the niche |
| Cubic polynomial | polynomial of degree 3 in *x* | 4 | a smooth response |
| Spline (10 df) | cubic B-spline in *x* | 10 | a smooth response |

Since the logarithm of a Gaussian niche, log *f*_*j*_ (*x*) = −(*x* −*θ*_*j*_)^2^/(2*b*^2^), is a polynomial of degree 2 in *x* with coefficients specific to the species, the quadratic basis is the correct specification. The categorical basis and the spline have the same number of coefficients and differ only in smoothness. As opposed to the categorical bases, we call *smooth bases* the ones in which the abiotic effect is a smooth function of the gradient (quadratic, cubic and the spline).

### 2.3. Scoring an estimated network

#### Ranking the pairs

Each estimator gives a path of networks, from empty to nearly full, as the penalty of the graphical lasso decreases. All the paths use the same grid, 25 penalties per decade from the smallest penalty that gives the empty network, and the same solver, that of PLNmodels; we checked that the depth of the paths does not affect the comparison. In practice a network is chosen along such a path by a criterion such as the BIC or the stability of the edges under resampling. We did not select a network on the path, so that the comparison does not depend on a selection rule: the pairs of species were ranked by the order in which they enter the path. Hence, a pair that enters at a larger penalty ranks higher, and two pairs that enter together are ordered by the size of their entry in the densest network. Competition and facilitation were scored separately, each with the sign it is expected to have. For competition, a pair counts when it enters with Ω_*jk*_ *>* 0; a pair that enters with the other sign, or never enters, is tied with the others of its kind at the bottom of the ranking. For the marginal readings, the pairs were ranked by the size of their residual correlation of the expected sign, negative for competition. An estimator is therefore rewarded only when it finds a pair *and* its sign.

#### Metrics

We consider two scores based on the ranking of pairs, one for each use. The first use is to rank all the pairs, and its score is the AUC (area under the ROC curve). It equals the probability that a pair drawn at random among the pairs that interact ranks above a pair drawn at random among those that do not, ties counting for one half: 0.5 for a random ranking and 1 for a perfect one. For competition, the pairs that interact are the competition pairs, and all the others count as non-interacting: the pairs with no interaction and the facilitation pairs. The reverse holds for facilitation. The AUC was computed exactly from the ranking of all the pairs, and not from the points of the path. The second use is to shortlist pairs, and its score is the *top-k precision*: the proportion of true interactions among the *k* best-ranked pairs, *k* being the number of true interactions of the sign considered in the network of the replicate (i.e., the precision of a network that would have the true density). The score is thus computed at a single value of *k*, set by each simulated network, and not aggregated over the values of *k*: on average *k* is about 7 for competition and 3 for facilitation at *D* = 10, and 29 and 14 at *D* = 50. At this value of *k*, the precision and the recall of the shortlist are equal. It measures how reliable the top of the ranking is, for a user who wants a small number of reliable candidate interactions.

#### Paired increments

A raw score does not measure what an estimator captures of the interactions, because the network is tied to the niches. Indeed, ranking the pairs by the proximity of their niche optima, without any data, already scores well above the chance level (Table S6). We therefore read every result as an *increment*: the score obtained on the simulation with interactions minus the score obtained on its paired no-interaction control (Section 2.1), both against the same network *W*. On the control the network does not act, and the score is what the estimator owes to the structure of the niches; the increment is what it owes to the interactions. Raw scores are reported next to the increments.

#### Rules of comparison

We set two rules before reading the results: first, each estimator is taken at its best abiotic basis, in each setting and for each metric, so that an estimator is never penalised for a basis that does not suit it (the results basis by basis are reported separately); second, partial associations are compared with partial associations, and the same for marginal ones. Each replicate was scored separately, and the increments were averaged over the 40 replicates of a setting. The difference between two estimators was computed setting by setting, and we report its mean over the settings of a mechanism, a 95% confidence interval, and the proportion of the settings in which it is positive.

### 2.4. Experimental design

Table 4 lists the experiments reported in the main text. The main design crosses the eight configurations, the three mechanisms and the strengths of the interactions (Table 1), and carries most of the results. The other two each vary a single factor, in a few settings. Three further experiments, reported in Supplementary Section S5, relax the simplifying assumptions of the simulators one at a time: a covariate measured with error or a second gradient that is not measured, other numbers of sites and species, and a dense interaction network of low rank.

**Table 4.** The experiments reported in the main text. The parameters take their default values unless they are the factor varied: *a* = 2 for the lottery, *κ* = 0.5 and *σ* = 0.5 for the Gompertz mechanism, *n* = 50 sites and *p* = 20 species. The supplementary experiments are listed in Table S7.

| Experiment | Settings | Factor varied | Estimators |
|---|---|---|---|
| Main design | 120: three mechanisms, eight configurations, all strengths | See Table 1 | levels 1 to 4, each with the five bases;<br>marginal readings of levels 1 and 3;<br>PLNPCA at ranks 2 and 5;<br>gllvm |
| Sampling effort | lottery and Gompertz; *b* ∈ {5, 10, 20} at *D* = 10, and *b* = 10 at *D* = 100 | spread of the effort *s* ∈ {0, 0.25, 0.5, 1} | levels 1 and 4, without correction and with each correction for effort |
| Abundance | Gompertz, *κ* ∈ {0.5, 0.9}; *b* = 10, *D* = 10 | *K* ∈ {3, 10, 30, 100, 1000} | levels 1, 3 and 4 |

#### Sampling effort

When the sites are sampled with unequal effort (Section 2.1), an estimator can ignore it, correct for it when the efforts *e*_*i*_ are known, or correct for it without knowing them. When the efforts are unknown, the data are *compositional*: they are still counts, but only the relative abundances of the species within a site are informative. PLNnetwork was fitted with no offset, with the offset *o*_*i*_ = log *e*_*i*_, and with the offset *o*_*i*_ = log ∑_*j*_ *Y*_*ij*_, the logarithm of the total count of the site, which requires no knowledge of the effort. The log-GGM was fitted on the counts as they are, on the counts divided by the effort, on the counts rescaled to a common total per site, and on the centred log-ratio of the counts (Aitchison, 1986), a transformation that has no counterpart in a model for counts, where the offset plays its role. To limit the number of fits, each was taken at the better of the cubic polynomial and the spline, the smooth bases that suit the two estimators best in the main design. At *D* = 100 every pair of species is eligible for an interaction, so that the network no longer follows niche proximity. This configuration is thus used as a the control.

#### Abundance

We underline that the different mechanisms are not matched for abundance: under the lottery, a site holds *K* = 100 individuals versus several hundred on average under the other two (Table S1). Under the Gompertz mechanism the constant *K* multiplies the expected counts and changes nothing else. Varying it shows whether an advantage of the count model comes from the small counts, where the transformation log(1 + *Y*) departs most from the logarithm of the expected count.

#### Software

All the computations were run in R, with PLNmodels 1.3.2 for the PLN, PLNPCA and PLNnetwork models and for the graphical lasso, gllvm 2.0.13 and Hmsc 3.3.7. A fit that failed, or that returned a likelihood that was not finite, was left out together with its paired fit on the control; this happened only with the basis that ignores the environment, for less than 5% of the fits of the count models (Supplementary Section S3).

## 3. Results

Competition and facilitation are reported side by side throughout, and the numbers are increments read as defined in Section 2.3: each estimator at its best abiotic basis, and a difference between two estimators given as its mean over the settings of a mechanism, followed in brackets by the proportion of the settings in which it is positive.

### 3.1. Effect of the configuration

How much there is to capture depends first on the room that the niches leave to the interactions (Table 5). With the narrowest niches (*b* = 2), the increment of PLNnetwork for the AUC of competition is small, and close to zero when the network does not follow niche proximity (*D* = 50). Under the lottery and the Gompertz mechanism it grows with the niche breadth, to about 0.3 at *D* = 10; under the Ricker mechanism the profile is flatter. At *D* = 50 the increments are lower everywhere. With narrow niches (*b* = 2) a site holds about two species (Table S1), and two species that interact seldom meet.

**Table 5.** Increment of PLNnetwork for competition, by niche breadth *b* and interaction threshold *D*: mean over the strengths of the interactions.

| metric | mechanism | *D* | *b* = 2 | *b* = 5 | *b* = 10 | *b* = 20 |
|---|---|---|---|---|---|---|
| AUC | Lottery | 10 | 0.08 | 0.17 | 0.27 | 0.25 |
|  |  | 50 | 0.02 | 0.05 | 0.09 | 0.16 |
|  | Gompertz | 10 | 0.07 | 0.20 | 0.28 | 0.33 |
|  |  | 50 | 0.01 | 0.07 | 0.14 | 0.20 |
|  | Ricker | 10 | 0.12 | 0.22 | 0.20 | 0.23 |
|  |  | 50 | 0.02 | 0.04 | 0.12 | 0.12 |
| top-*k* precision | Lottery | 10 | 0.14 | 0.26 | 0.27 | 0.23 |
|  |  | 50 | 0.03 | 0.07 | 0.12 | 0.14 |
|  | Gompertz | 10 | 0.11 | 0.23 | 0.34 | 0.40 |
|  |  | 50 | 0.03 | 0.10 | 0.17 | 0.23 |
|  | Ricker | 10 | 0.21 | 0.17 | 0.17 | 0.21 |
|  |  | 50 | 0.03 | 0.08 | 0.11 | 0.14 |

### 3.2. Partial against marginal associations

We compared the partial and the marginal readings of the network in a setting in which nothing else changes: the sparse precision matrix of PLNnetwork against the correlations read in the covariance of the same PLN fit, on the same basis and the same replicate (Figure 1). For the top-*k* precision, the partial reading is ahead by 0.04 to 0.07 under the three mechanisms, for competition and for facilitation, in 72 to 82% of the cases. Over the whole ranking the gain is small, at most 0.02 for the AUC.

**Figure 1.**
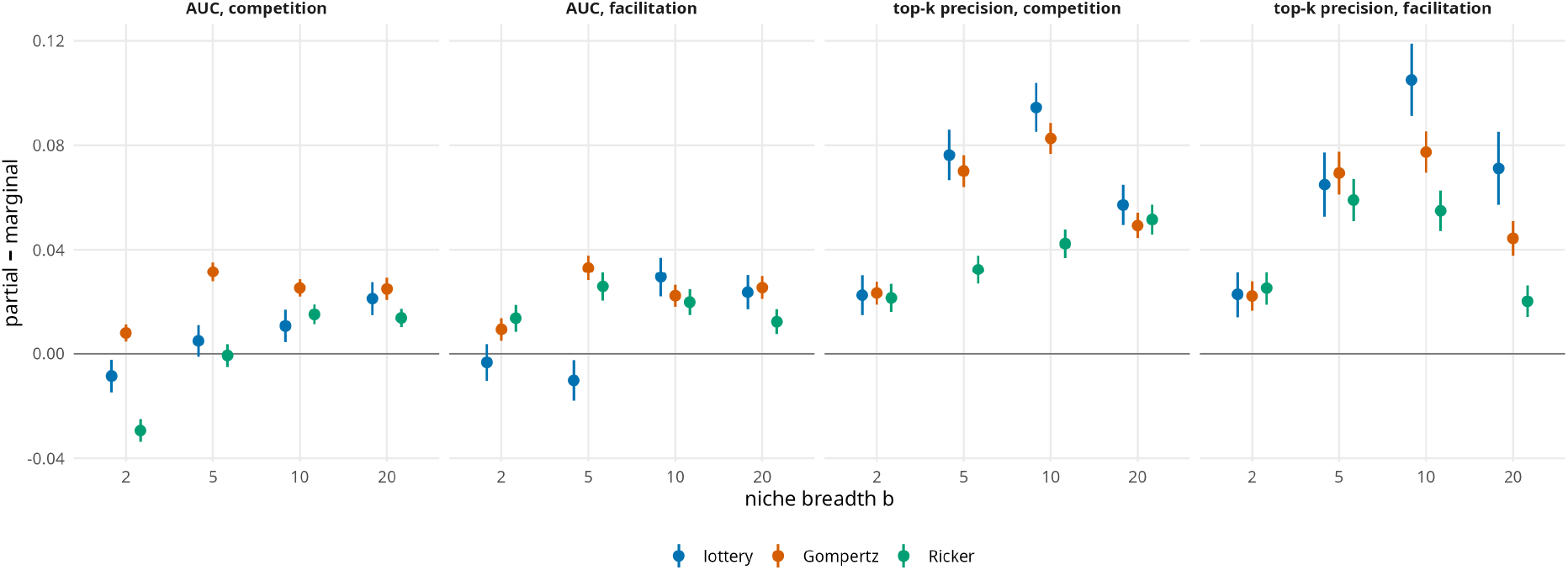
Partial against marginal associations. Increment of PLNnetwork minus increment of the correlations read in the covariance of the same PLN fit, on the same basis and replicate, against the niche breadth *b*; the five bases and all the strengths are pooled. Above zero, the partial reading does better. Bars: 95% confidence intervals.

The gain depends on the configuration: for the AUC it grows with the niche breadth, and it is slightly negative for competition with the narrowest niches under the lottery and the Ricker mechanism; for the top-*k* precision it is largest at intermediate niche breadths (*b* = 5 and *b* = 10). Reading partial associations instead of marginal ones therefore matters most to a user who wants a short list of reliable pairs.

### 3.3. Comparison of the four levels of integration

Figure 2 compares the four levels of integration. We first set the integrated JSDM, PLNnet-work (level 4), against the log-GGM (level 1). Under the Gompertz mechanism, PLNnetwork is ahead on the four metrics, by 0.02 for the AUC and about 0.04 for the top-*k* precision, in 85 to 92% of the settings. Under the lottery, it keeps one advantage, the top-*k* precision of competition (+0.04, 92%); for the two AUCs and for the top-*k* precision of facilitation, the log-GGM does as well or better. Under the Ricker mechanism, the log-GGM does as well or better on the four metrics.

**Figure 2.**
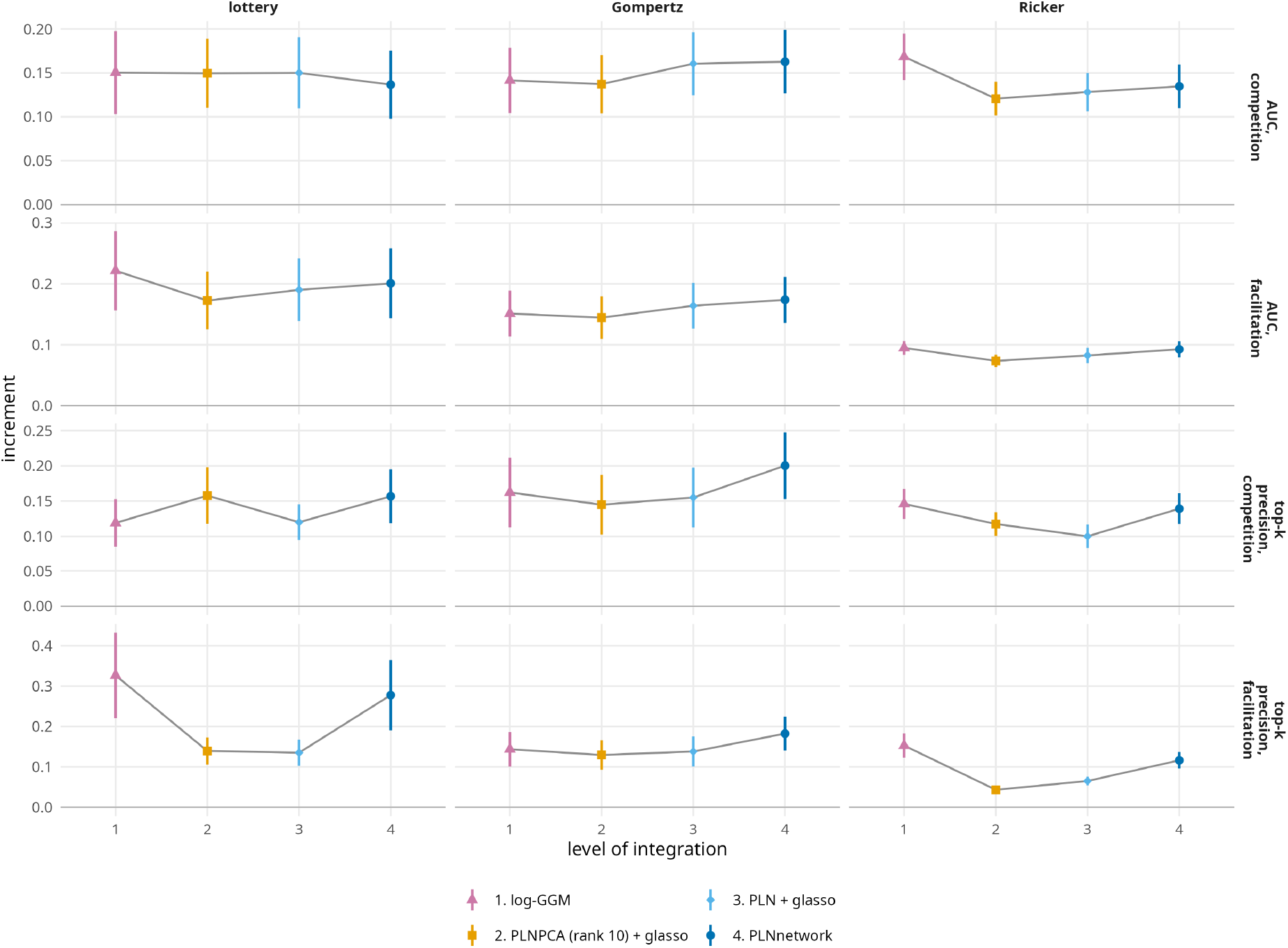
The four levels of integration. Mean increment of each estimator at its best basis, by mechanism (columns) and metric (rows). One observation per setting (24 under the lottery, 48 under the Gompertz and Ricker mechanisms); bars: 95% confidence intervals over the settings. The raw scores are in Figure S9.

Since the Ricker mechanism differs from the Gompertz mechanism by log-linearity alone, the integrated JSDM is ahead over the whole ranking when the mechanism is log-linear, that is when the network is written in the precision matrix of its latent layer, and not otherwise. The comparison of the lottery and the Gompertz mechanism on the same communities and the same networks agrees (Supplementary Section S9, Figure S11): from the first to the second, the advantage of PLNnetwork over the log-GGM rises for the two AUCs and for the top-*k* precision of facilitation, and does not change for the top-*k* precision of competition, for which PLNnetwork was already ahead under the lottery. Under the Gompertz mechanism, the advantage is largest with narrow niches and a network that follows niche proximity (Figure S10).

Two remarks temper these comparisons. First, the gain is not incremental from one level to the next: levels 2 and 3 are often below level 1, and under the Ricker mechanism the three estimators built on a count model are all below the log-GGM for the AUC of competition. Second, the scores themselves are modest. On the simulations with interactions, the raw top-*k* precision of levels 1 and 4 lies between 0.19 and 0.41 (Figure S9).

### 3.4. Sources of the advantage of the integrated JSDM

The steps between levels separate what the integrated JSDM owes to its model for counts, to the rank of its covariance and to the integration (Figure 3).

**Figure 3.**
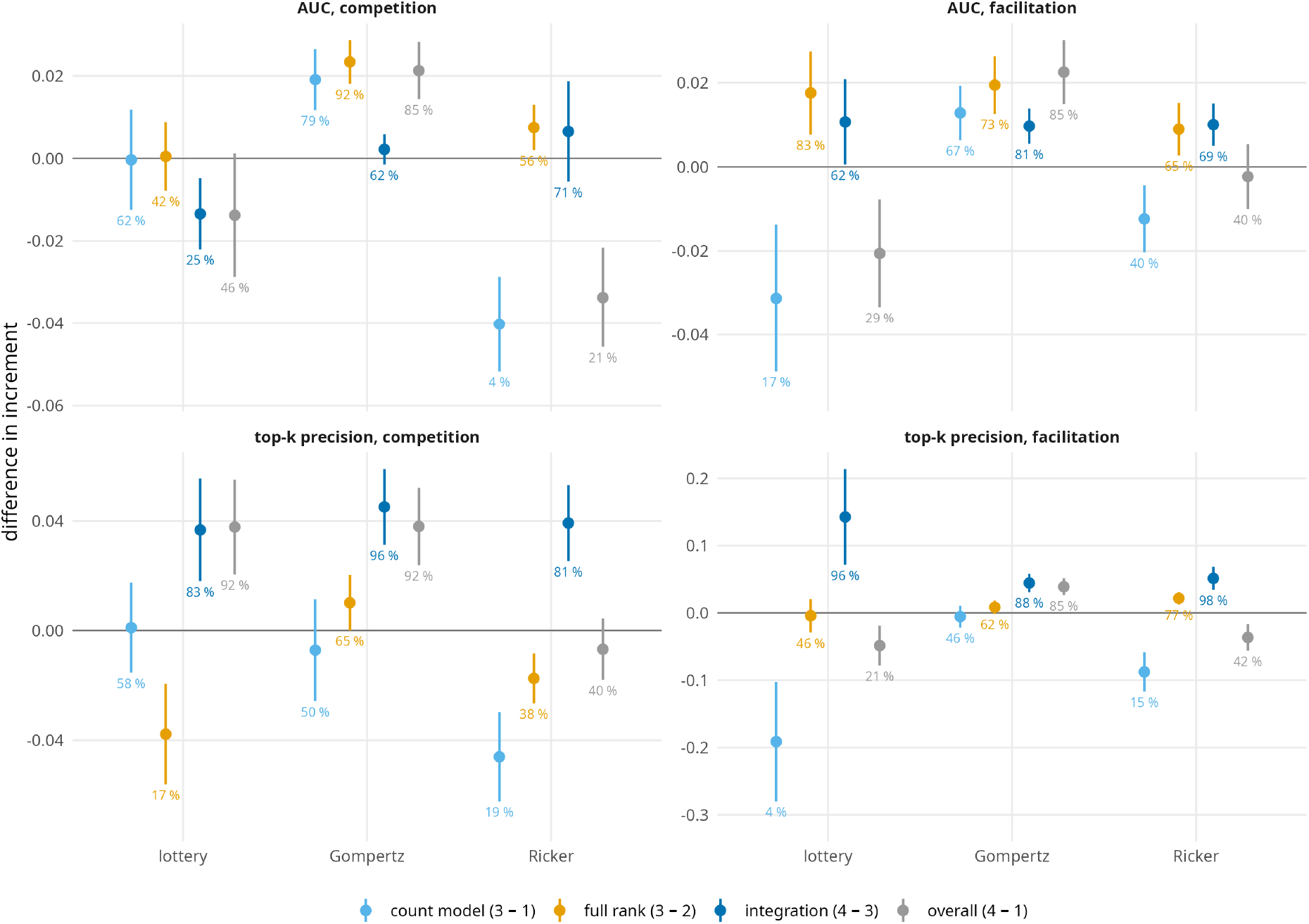
Origin of the gain brought by each step between two levels: difference of increments, setting by setting, each estimator at its best basis. Count model: level 3 minus level 1; rank: level 3 minus level 2; integration: level 4 minus level 3; total: level 4 minus level 1. Mean and 95% confidence interval over the settings; the percentages are the proportions of the settings in which the step is positive.

*The count model* (from level 1 to level 3) acts on the whole ranking, and under the log-linear mechanism only: the AUC of competition gains about 0.02 under the Gompertz mechanism, nothing under the lottery, and loses about 0.04 under the Ricker mechanism. At the top of the ranking it brings nothing for competition, and it costs for facilitation under the lottery and the Ricker mechanism.

This gain comes from the small counts. When the constant *K* of the Gompertz mechanism is varied (Table S15), the advantage of PLNnetwork over the log-GGM for the AUC of competition is largest at *K* = 10 and shrinks as the counts grow; at *K* = 3 there is almost nothing left to capture for any estimator.

*The integration* (from level 3 to level 4) acts at the top of the ranking, under the three mechanisms: the top-*k* precision gains about 0.04 for competition, and as much or more for facilitation, in more than 80% of the settings, while the AUC hardly moves. A graphical lasso applied to the covariance of a fitted JSDM therefore does not replace an estimator that looks for the sparse network while fitting the model, when the aim is a short list of pairs.

*The rank* (from level 2 to level 3) is a constraint that costs. Every latent factor added to PLNPCA helps (Figure 4). At rank 10, half the number of species, PLNPCA is close to the model of full rank. At rank 2, the default of gllvm, it is behind the log-GGM everywhere, with about half to two thirds of its increment. gllvm with its two latent variables gives the same results as PLNPCA at rank 2 (Supplementary Section S6).

**Figure 4.**
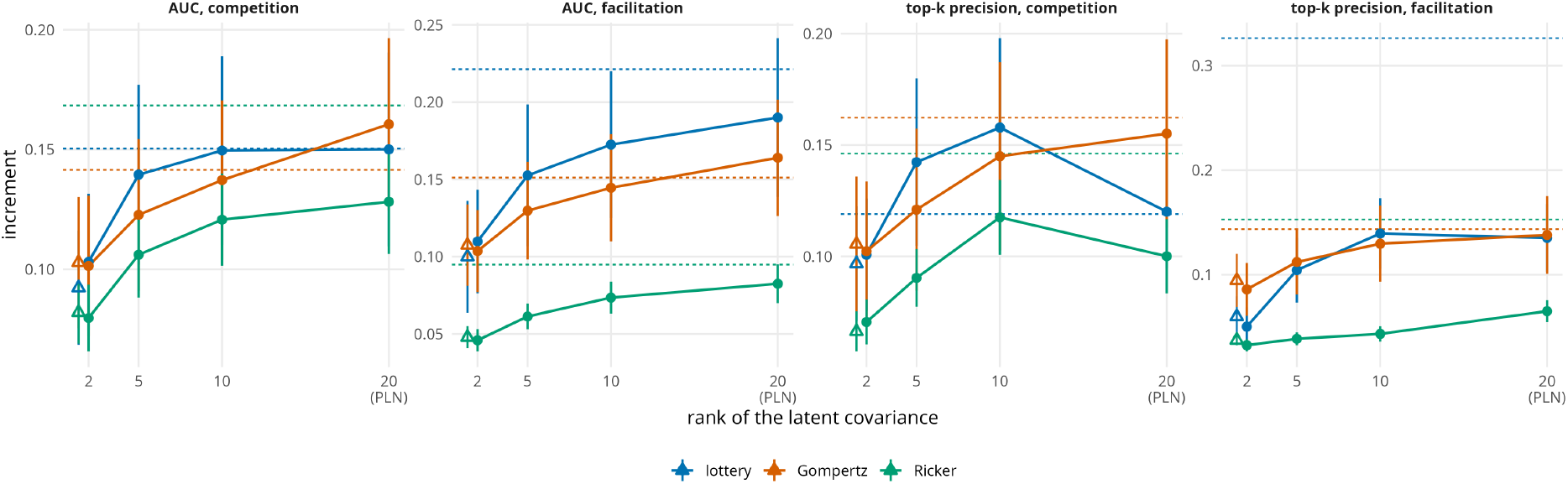
The rank of the latent covariance. Mean increment of PLNPCA followed by the graphical lasso at ranks 2, 5 and 10, and of PLN followed by the graphical lasso (full rank, 20 species), each at its best basis. Open triangle: gllvm with 2 latent variables, followed by the graphical lasso. Dashed line: the log-GGM.

### 3.5. Unequal sampling effort and compositional data

An unequal sampling effort that is not corrected for creates a network signal where there is no interaction (Figure 5). A site that is sampled more yields more individuals of all the species it holds, so that the species that co-occur vary together; and at *D* = 10 the species that interact are among those that co-occur. The paired no-interaction control reveals this signal, since its score should not depend on the effort. Without correction, the raw AUC of the control drifts as the spread of the effort grows, upwards for facilitation with PLNnetwork and downwards for competition with the log-GGM.

**Figure 5.**
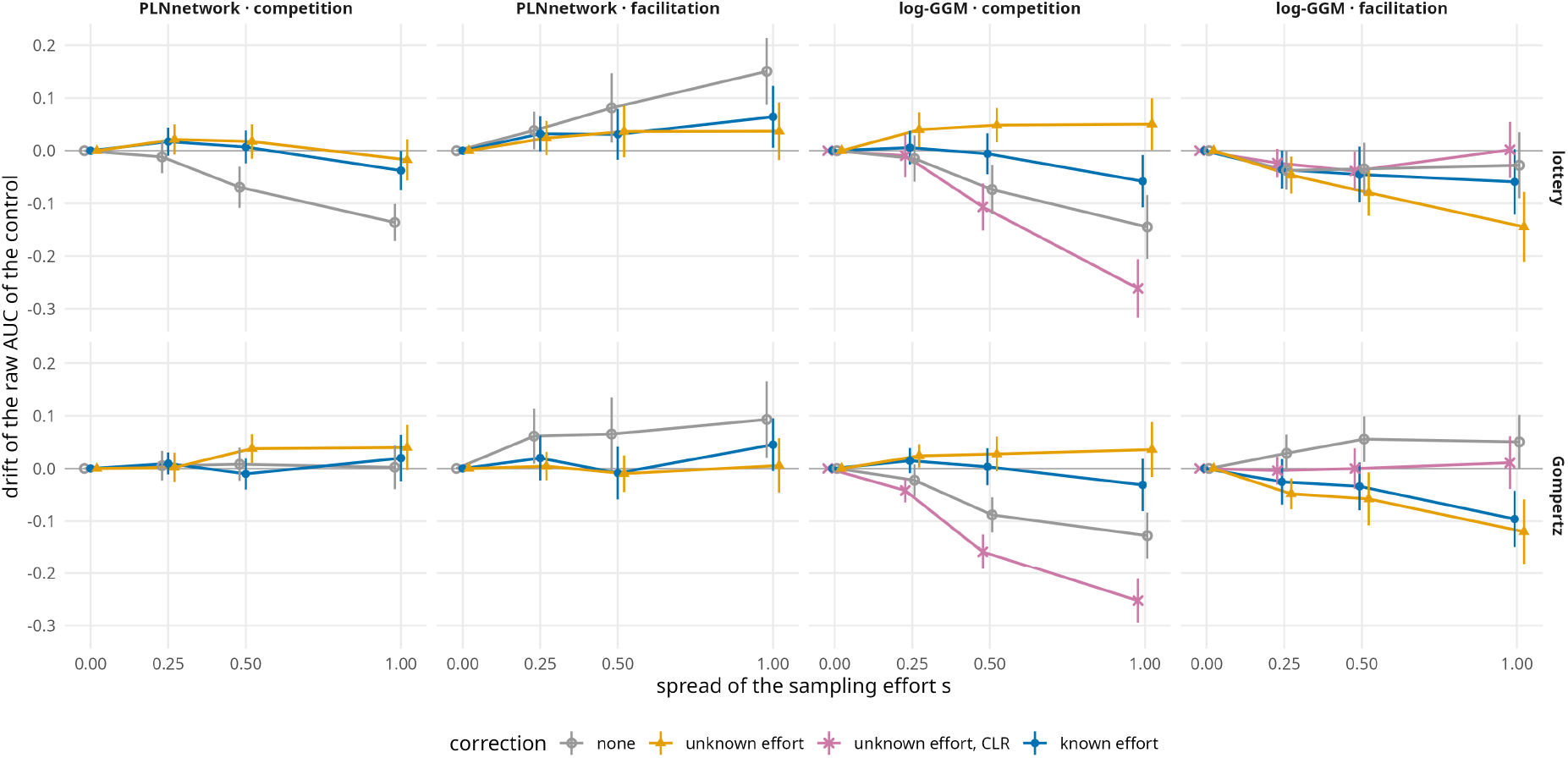
Unequal sampling effort creates a spurious signal. Drift of the raw AUC of the paired no-interaction control (its value minus its value at *s* = 0) against the spread *s* of the effort, at *b* = 10 and *D* = 10, by mechanism (rows), estimator and type of interaction (columns). Corrections: none; unknown effort (PLNnetwork: offset equal to the logarithm of the total count of the site; log-GGM: counts rescaled to a common total); unknown effort, centred log-ratio (log-GGM only); known effort (PLNnetwork: offset log *e*_*i*_; logGGM: counts divided by the effort). Without interactions the drift should be zero. Bars: 95% confidence intervals over the 40 replicates.

With either offset, from the known effort or from the total count of the site, the control of PLNnetwork drifts little. For the log-GGM, dividing the counts by a known effort limits the drift, whereas rescaling the counts to a common total does not stop it, and the centred log-ratio makes it larger for competition. The increments themselves are given against the spread in Figure S12.

The raw scores of the two arms (Figure S13) show why a raw score must be read against its control. Under the Gompertz mechanism, the raw AUC of facilitation of the log-GGM is high, but so is that of its control: what it measures is the niche structure left in its residual associations, which facilitation pairs share since their niches are close. In increments, PLNnetwork captures more of facilitation than the log-GGM under this mechanism.

What a correction is worth then depends on the mechanism (Figure 6, at *s* = 0.5). Under the Gompertz mechanism, the count model with an offset is the best reading, with the known effort as with the total count of the site, which requires no knowledge of the effort. Under the lottery, no correction helps PLNnetwork, and the log-GGM on the counts divided by the effort does better. The offset is the right correction when the effort multiplies the mean of a Poisson log-normal count, which is the case under the Gompertz mechanism and not in the closed community of the lottery, where the number of individuals per site is fixed. At *D* = 100, where the network no longer follows the niches, all the readings give about the same increment.

**Figure 6.**
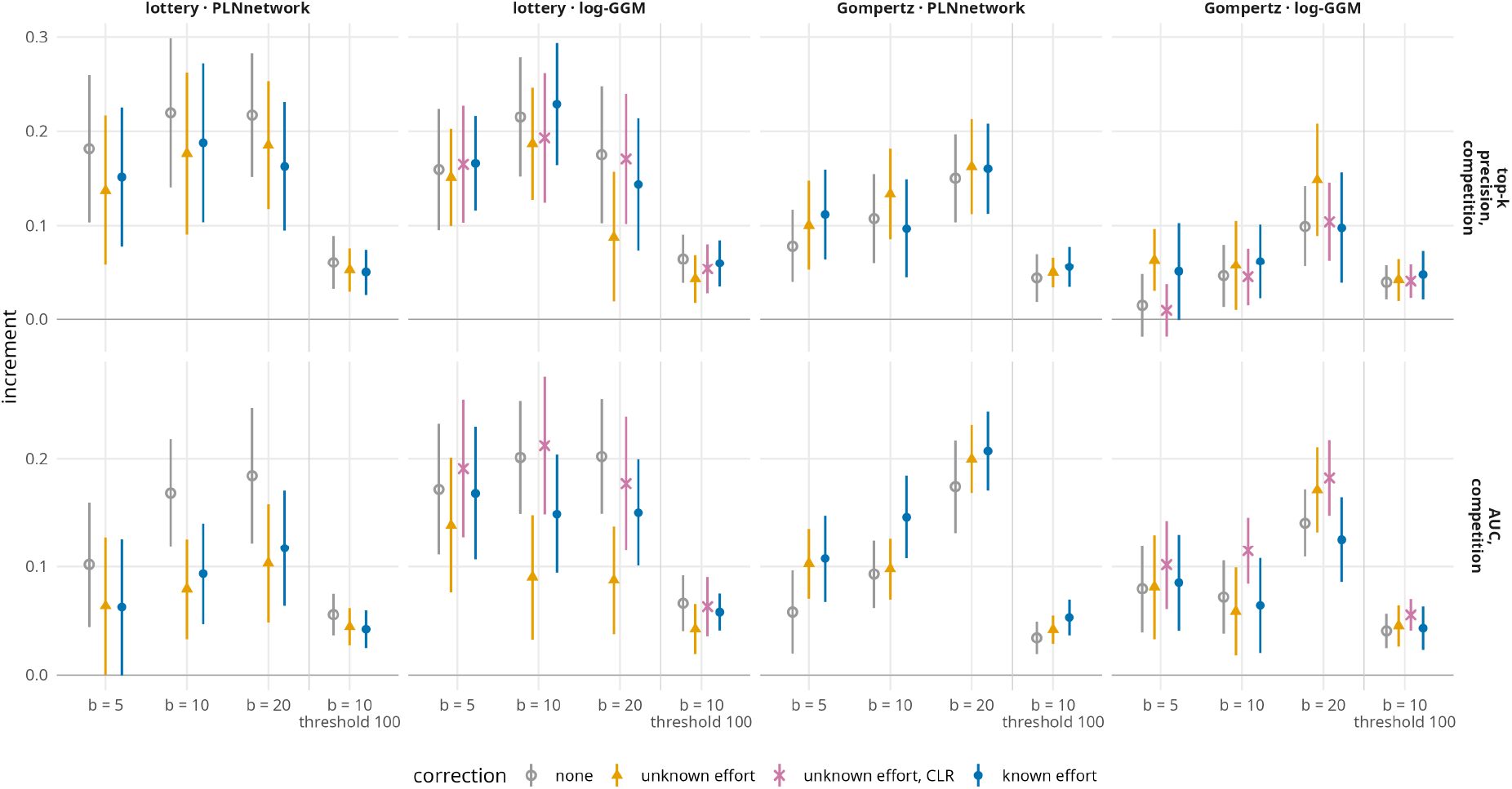
Corrections for effort across configurations: increment at a spread *s* = 0.5, each reading at its best basis, by mechanism and estimator (columns) and metric (rows); corrections as in Figure 5. In the last configuration (*D* = 100) the network does not follow niche proximity. Bars: 95% confidence intervals over the 40 replicates.

### 3.6. Choice of the abiotic basis

The five bases show that there is no single appropriate basis, and what the choice depends on (Figure 7).

**Figure 7.**
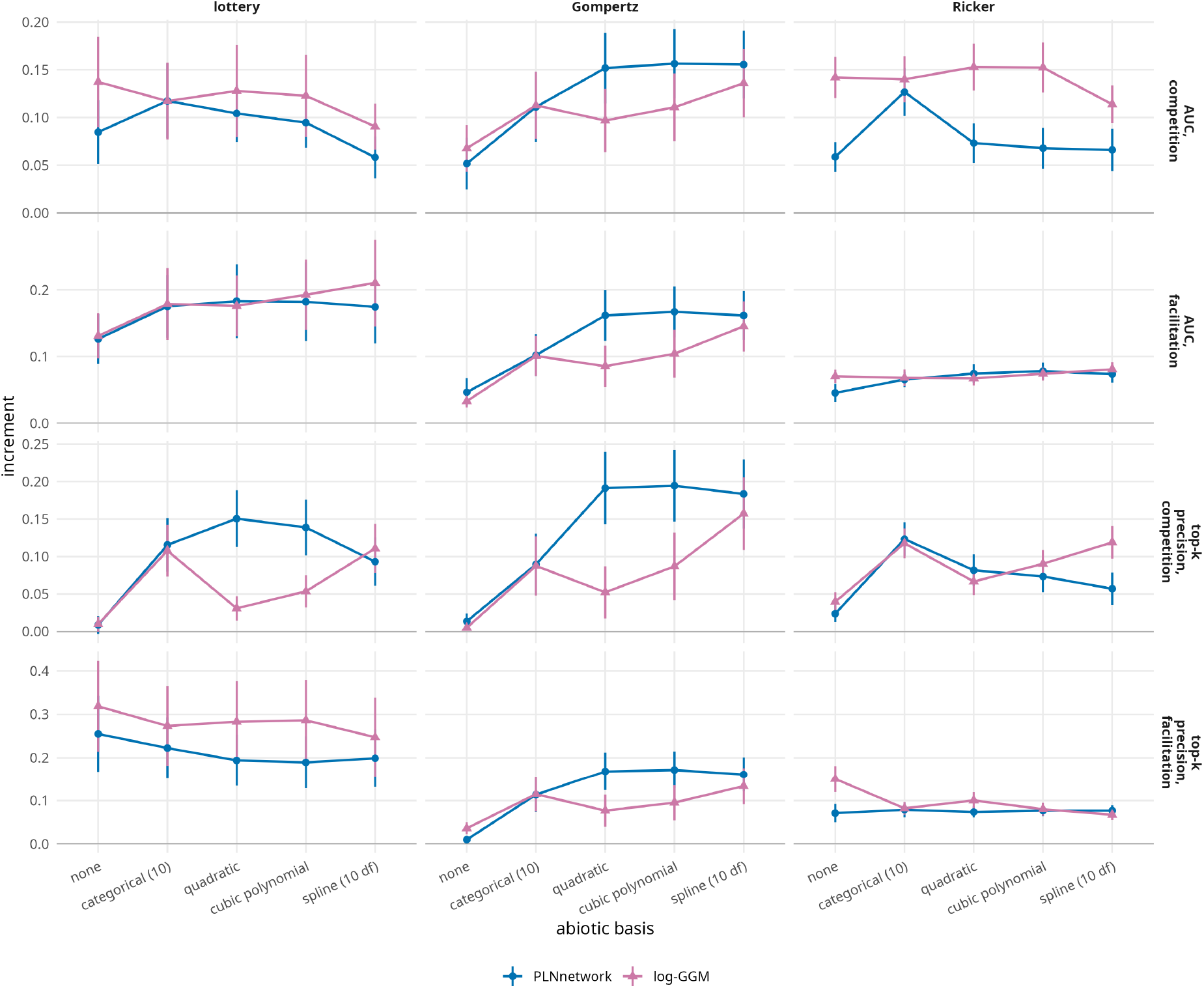
Each estimator has its abiotic basis. Mean increment of PLNnetwork and of the log-GGM for each of the five bases, over all the configurations and strengths; bars: 95% confidence intervals.

#### Dependence on the estimator

Under the Gompertz mechanism, PLNnetwork draws as much from any smooth basis, and less from the categorical basis or from none. The log-GGM comes close only with the spline: the logarithm of one plus a count that follows a Gaussian niche, flattened by the zeros, is not a quadratic function of the gradient. This is why each estimator is taken at its own best basis.

#### Dependence on the mechanism

Under the lottery and the Ricker mechanism, the categorical basis is the best one for PLNnetwork and the AUC of competition. Moreover, under the lottery the top-*k* precision of facilitation is highest with no covariate at all, for both estimators. Competition and facilitation thus call for different bases under the lottery; under the Gompertz mechanism this opposition disappears, the smooth bases being the best for both types of interaction. The basis that serves an estimator best is not the one that recovers the niches best (Supplementary Section S9).

The basis is not the only condition. The supplementary experiments (Supplementary Section S5) show that the advantage of the count model under the Gompertz mechanism also requires a covariate measured almost without error, and no gradient that structures the community without being measured.

## 4. Discussion

We have introduced a simulation experiment that falls into the “virtual ecologist” approach (Zurell et al., 2010) to question the relationship between statistical patterns and the processes defined by biotic interactions in the context of JSDMs. In this section, we start by discussing to what extent competition and facilitation are captured in partial associations. We then turn to what an integrated JSDM for counts brings beyond a simpler estimator, to the difference between competition and facilitation, and to the way abiotic effects are accounted for. Finally, we discuss the limits of this experiment and the perspectives it opens.

### 4.1. Capture of biotic interactions by partial associations

The idea that residual (marginal or partial) correlations as inferred by JSDMs may relate to biotic interactions is based on the intuition that species in competition (resp. facilitation) cooccurr less (resp. more) than species with no interactions (Morales-Castilla et al., 2015; Freilich et al., 2018), although there is no clear proof that co-occurrences can inform biotic interaction networks in general (Freilich et al., 2018; Poggiato et al., 2021, see). Thus, in our simulations, we expect JSDM-inferred association networks to match, at least partially, the simulated biotic interactions, with negative residual correlations observed where competition occurs and positive ones for facilitation. This expectation is exact, to first order, under one of our three mechanisms only, the Gompertz mechanism, in which the network is written in the precision matrix of the latent layer of a Poisson log-normal model. Under the lottery and the Ricker mechanism, it rests only on the intuition above.

The results confirm the expectation and bound it. Competition and facilitation are captured under the three mechanisms, in the sense that every estimator scores higher, on average, on the simulation with interactions than on its paired no-interaction control. They are captured better in partial than in marginal associations, which agrees with what was established with other models and other simulators (Harris, 2016; Popovic et al., 2019), and adds one precision: on the same fit, the gain is small over the whole ranking and clear among the pairs ranked first. However, two things limit what is captured. The first is the configuration of the niches: when species that interact seldom meet, as with narrow niches, there is very little to capture for any estimator. The second is the level of the scores. A raw AUC cannot be read on its own, since ranking the pairs by the proximity of their niches already gives a nearly perfect AUC when the interacting species have close niches (Table S6). Once the part that a score owes to the niches is removed, what remains is modest, and of the pairs ranked first by the best estimators, one in five to two in five are true interactions. This confirms the difficulties inherent to the task of relating (statistical) patterns to biotic mechanisms: while competition appears as a legitimate hypothesis to explain a negative residual association, it requires further experimenting to be confirmed.

### 4.2. Contribution of an integrated JSDM for counts

The four estimators that we compared are four ways of doing the same thing: separating what the environment explains from the dependence between species, and reading a sparse network in the latter. The simplest of them, the Gaussian graphical model on log-transformed counts, is itself an approximate JSDM, in which a transformation stands for the model for counts. The comparison therefore does not oppose JSDMs to something else: it shows which ingredient of an integrated JSDM for counts helps, and when.

The model for counts helps when it is the right model. Under the Gompertz mechanism the logarithm of the abundances is Gaussian and the network sits in its precision matrix: the Poisson log-normal model is correctly specified, and it ranks the pairs better than the transformation does, all the more as the counts are small or the sampling effort unequal, that is where the transformation log(1 + *Y*) departs most from the logarithm of the expected count. Under the lottery and the Ricker mechanism the network is not the precision matrix of any Gaussian layer. Both estimators are then misspecified, and the simpler one loses nothing. The Ricker mechanism shows that this is a matter of log-linearity and not of the nature of the data, since it shares with the Gompertz mechanism everything but log-linearity. An analyst does not know which mechanism produced the data. What these results say is that the model for counts cannot be counted on to improve the ranking of all the pairs, and that it is the safer choice when counts are small and sampling effort varies.

This is not a weakness of the Poisson distribution of the Poisson log-normal model. The Poisson layer only describes the sampling of the counts given the latent vector. What the lottery and the Ricker mechanism do not satisfy is the Gaussian latent layer on the log scale, in which the network is read and which the most widely used JSDMs share, whatever their emission distribution. We checked it with gllvm, fitted with a Poisson or a negative binomial distribution, at the same number of latent variables and with the same graphical lasso (Supplementary Section S6): under the Gompertz and the Ricker mechanisms the two distributions give similar increments, and under the lottery the negative binomial does much worse. In no case does another emission distribution bring a latent-factor JSDM to the level of the log-GGM under the lottery or the Ricker mechanism.

The integration helps for another use, and under the three mechanisms. A sparse precision matrix estimated together with the model is more reliable among the pairs ranked first than one obtained by applying the graphical lasso to a covariance fitted without penalty. This concerns a common practice, which consists in inverting, or applying the graphical lasso to, the covariance reported by a JSDM: this gives partial associations, but their best-ranked pairs are less reliable than those of an estimator that looks for the sparse network while fitting the model.

The rank of the latent covariance is the third ingredient, and the one that separates the latentfactor JSDMs from the others. With a covariance of rank two, the associations between twenty species are carried by two factors only, and the network read in it is behind that of the simple Gaussian model on every metric. This is an effect of the rank and not of a piece of software: gllvm, in its Poisson version, and a Poisson log-normal model of the same rank give the same results, and each factor added helps. With two latent variables, a common default, a latent-factor JSDM should not be expected to capture interactions between pairs of species.

The four estimators obtain a sparse network through an *l*_1_ penalty, and none of them comes with a test or a confidence interval for an association. This is not specific to the Poisson lognormal models: as for the lasso in regression, inference after an *l*_1_-penalised estimation is a question of post-selection inference, which remains open for graphical models of counts. In practice a network is chosen along the path of penalties by a criterion, such as the stability of the edges under resampling, and an interaction cannot be defined by a significantly positive or negative association. This is consistent with what the results support: an estimated network is a ranking of pairs, whose top is a short list of candidates, and not a set of tested interactions.

### 4.3. Competition and facilitation

Competition and facilitation can call for opposite treatments of the abiotic effect, which can be explained by the joint effect of the abiotic and biotic filters. For competition the two are antagonistic: species can only be in competition if their niches are close enough so that they still have more chances of being found together than species with distant niches that would not be in competition. For facilitation they are synergistic: two species involved in facilitation have two reasons to be found together, their niches are close and they favor each other’s presence.

Therefore, it is likely that when accounting for the environmental effect, facilitation is “absorbed”, in the abiotic filters through the “mean effects” of the underlying generalized linear model. However, when the covariate is not included, both the abiotic and biotic filters appear in the positive residual correlations, with the strongest positive correlations observed for species with facilitation interactions. As a side effect, it is likely that species with very close niches are also more likely to be mistakenly identified as being involved in facilitation interaction. The paired no-interaction control now measures this side effect, since it has the same niches and no interaction: this is the part of a score that the increment removes.

The basis acts on the network only through what it leaves in the residuals: the abiotic coefficients absorb the part of the abundances that varies along the gradient in the way the basis allows, including the part due to the interactions when they change the mean abundance of a species along the gradient. The explanation above therefore holds under the lottery, in which a facilitator raises the chance that a species occupies a site: there the top-*k* precision of facilitation is highest with no covariate at all. It does not hold under the Gompertz mechanism, in which the interactions shape the fluctuations of the abundances around the level set by the environment, and not that level (Supplementary Section S4, Table S5): the abiotic effect has nothing to absorb.

These results underline that a JSDM’s ability to disentangle different mechanisms (biotic, abiotic…) through the statistical patterns it unveils strongly depend on the joint dynamics of these mechanisms and more specifically, whether their effects are synergestic or antagonistic, an idea that Poggiato et al. (2021) also underlined. They add that it depends on the mechanism by which the interactions act, which is not known for real data.

### 4.4. Accounting for abiotic effects

When accounting for covariates, several approaches can be used. The most natural one, used in several JSDMs, including the PLN-network model, is to resort to a generalized linear model (GLM) and directly plug the covariates in it, with no or little transformation. However, in this simulation framework, this does not appear as the most relevant approach. Indeed the (virtual) environmental covariate covers a large gradient and the environmental filter is based on the distance between a species’ niche and the local environment. A GLM with no covariate transformation then does not appear appropriate as it cannot model the proximity between an environment and a niche but only general trends. Cutting the gradient into classes is one way to meet this difficulty. We compared five ways of entering the gradient, and none is the appropriate one in general. The basis that serves an estimator best depends on the estimator, a Poisson log-normal model drawing as much from any smooth basis when the transformed counts need a flexible one, and on the mechanism, as discussed above. Finally, it is not the basis that recovers the niches best: the accuracy of the abiotic estimation does not predict the quality of the network (Supplementary Section S9, Table S16). In practice we see no better course than to compare several bases and to trust what is stable across them.

A suitable basis does not make the abiotic and the biotic effects identifiable. A model separates them only through what the covariates explain, and whatever the covariates leave of the environment ends up in the residual associations. We measured three forms of this. An error of a quarter of a niche breadth on the covariate is enough to cancel the advantage of the model for counts under the Gompertz mechanism, and to leave enough niche structure in the associations for the raw ranking to be worse than a random one. A gradient that structures the community and is not measured removes most of what any estimator captures. Finally, an unequal sampling effort that is not corrected for creates a network signal where there is no interaction; an offset corrects it when the counts are samples of an open community, and not in the closed community of the lottery. In each case the control without interactions is what reveals the problem, and no such control exists for real data.

### 4.5. Limits and perspectives

Although this experiment shows that JSDMs can partially capture competition and facilitation provided they appropriately account for abiotic effects, there are limits to these results. First, although the VirtualCom model is designed to simulate realistic ecological processes (Münkemüller and Gallien, 2015), it remains a stereotypical framework, with interactions modelled in accordance with the same intuition as the one that underpins the relationship between biotic interactions and JSDM-inferred associations. Its lottery reproduces the consequence that an interaction is expected to have on the composition of a community, and not a process acting on populations. The Gompertz and Ricker mechanisms add such processes, but the three simulators keep the same simplifications: one environmental gradient, Gaussian niches of equal breadth, no dispersal between sites, and symmetric interactions between pairs of species, all of the same intensity. Real communities depart from each of them, and the experiment in which a second gradient is left unmeasured shows how much harder the task then becomes.

Asymmetric interactions are not studied here. An association is symmetric by construction, which is often named as a limit of JSDMs for the study of interactions (Dormann et al., 2018), and other interactions could lead to even more complex effects and statistical patterns, such as parasitism or trophic interactions. Interactions that affect one species only, such as commensalism or amensalism, should leave a pattern of the same kind as facilitation or competition, whereas for interactions that benefit one species at the expense of the other, such as predation or parasitism, the expected sign of the association is unclear.

The comparison has its own limits. It bears on communities of 20 species at 50 sites; with 40 species the advantage of the model for counts shrinks. It includes neither the copula graphical models (Popovic et al., 2019), the closest competitors of the Poisson log-normal models, nor a JSDM with a sparse covariance, and it says nothing of Hmsc, whose sampler did not converge in our hands. It scores networks and not predictions of abundance. The zeros of our simulations come from the niches and from sampling, never from a process that would exclude a species from a site whatever its niche, so that models for zero-inflated counts (Tous et al., 2026) were not considered.

Moreover, it is unclear to what extent the species distribution data that our simulators output can be compared to any real species distribution data set. When using a JSDM for a specific species distribution data analysis, it could be useful to re-use this framework and adapt the simulation parameters to obtain configurations that are closer to the considered dataset. The paired no-interaction control and the two uses of a ranking make a protocol that applies to any estimator of an association network, provided a few pitfalls are avoided (Supplementary Section S7), and the simulators can be given the numbers of sites and species, the abundances and the sampling effort of a data set at hand. What such an exercise can tell an analyst is how much a network estimated from data of that kind would retain of interactions of a given kind. It cannot tell which associations of the real network are interactions.

## 5. Conclusion

We have introduced an original virtual framework to test the relationship between statistical patterns identified by JSDMs and the processes induced by two symmetric biotic interactions, competition and facilitation. It rests on three interaction mechanisms, on estimators that differ by one ingredient at a time, on two uses of an estimated network, and on a paired control without interactions that removes what a score owes to the niches.

Whether a JSDM captures these interactions depends on conditions that we can now state. The configuration of the niches comes first: when interacting species seldom meet, no estimator captures much. Reading partial associations instead of marginal ones helps, mostly for the pairs ranked first. Beyond that, the two ingredients of an integrated JSDM for counts do not serve the same use. Its model for counts improves the ranking of all the pairs when the mechanism that generated the data is log-linear, and all the more as the counts are small or the sampling effort unequal; under the two other mechanisms a Gaussian graphical model on log-transformed counts, fitted in a few hundredths of a second, does as well. The estimation of the sparse network together with the model improves the short list of best-ranked pairs under the three mechanisms. A latent-factor model with two factors, a common default, is behind all of these. For the analyst of real data, who knows neither the mechanism nor the interactions, three recommendations follow. Read partial associations, and read them as a short list of candidate pairs rather than as a network. Compare several abiotic bases, since the appropriate one depends on the estimator and on the mechanism, and correct for sampling effort with an offset when counts are samples of an open community. Finally, do not expect a network estimated from one imperfectly measured gradient to separate interactions from what the covariates leave unexplained. These relationships should be analysed with care, paying attention to the inclusion and modelling of abiotic filters.

## Acknowledgements

We thank Wilfried Thuiller for his comments on the first version of the manuscript, which helped us state the questions from the point of view of an ecologist, and for sharing his knowledge of joint species distribution models; Laura Pollock for the idea of a distance threshold between niche optima, beyond which two species do not interact; and the recommender, Olivier Gimenez, and the two reviewers, whose many comments led us to improve the article substantially.

## Use of generative artificial intelligence

Jeanne Tous and Julien Chiquet designed the study. Jeanne Tous wrote the first version of the article, with minor revisions by Julien Chiquet. We rethought the second version following the many suggestions of the recommender and of the reviewers. For this version, we revised the text with the help of a generative artificial intelligence tool, Claude Code (Anthropic), English not being our native language. We also used it to reorganise and correct the code, and to set up the new simulation campaign; the README of the code repository details these uses. We checked the whole text and code, and take responsibility for them.

## Fundings

The authors declare that they have received no specific funding for this study.

## Conflict of interest disclosure

The authors declare that they comply with the PCI rule of having no financial conflicts of interest in relation to the content of the article.

## Data, script, code, and supplementary information availability

The Supplementary material (Sections S1 to S9: the simulators, the simulated data, the models compared, checks, supplementary experiments and additional results) is provided as a separate document with this submission. The code of the simulations and of the analyses is available at https://github.com/jchiquet/jsdm_biotic_interactions; its README describes how to reproduce every number and figure of the article. The results of the simulations, from which every number, table and figure is computed, are deposited on Zenodo (doi:10.5281/zenodo.24001750); the simulated data themselves are regenerated by the code. The Supplementary material and a versioned copy of the code will be deposited in an open repository, with a DOI, if the article is recommended.

## Notes

https://doi.org/10.5281/zenodo.20273114

### Competing Interest Statement

The authors have declared no competing interest.

### Summary of Updates

We rebuilt the simulation design and the comparison of the estimators. Such a revision has changed the message of the article in depth, from a comparison of three JSDMs to a study of the conditions under which a JSDM captures biotic interactions. The numbers have changed as well, mainly because the simulation protocol and the worlds it explores were rebuilt entirely: the communities are now simulated under three interaction mechanisms instead of one, over eight configurations of the niches instead of the single one of the submitted version, which turned out to leave little to capture, and every score is read against a paired no-interaction control instead of the chance level, since a raw score mostly reflected the environment. While rebuilding the protocol, we also corrected two defects in our scoring. The results of the submitted version are therefore superseded, and several of its conclusions are qualified; we say so at the outset, since many of our responses depend on it.

