## Supplementary material for "When can a joint species distribution model capture biotic interactions? A simulation study across interaction mechanisms and levels of model integration"

for the article

This document gathers what the main text refers to as the Supplementary material. Its sections, figures, tables and pages are numbered with an S; a number without an S is that of a section, a figure, a table or an equation of the main text. The code of the simulations and of the analyses is available at `https://github.com/jchiquet/jsdm_biotic_interactions`, and the results of the simulations on Zenodo (doi:10.5281/zenodo.24001750).

#### Contents

**S1 The lottery for space, step by step** S2

**S2 The simulated data** S2

**S3 The models compared** S4

**S4 Checks on the simulators and on the scoring** S8

**S5 Supplementary experiments** S11

**S6 gllvm and Hmsc** S16

**S7 Pitfalls in scoring an estimated network** S18

**S8 Run times** S18

**S9 Additional results of the main design** S18

### S1 The lottery for space, step by step

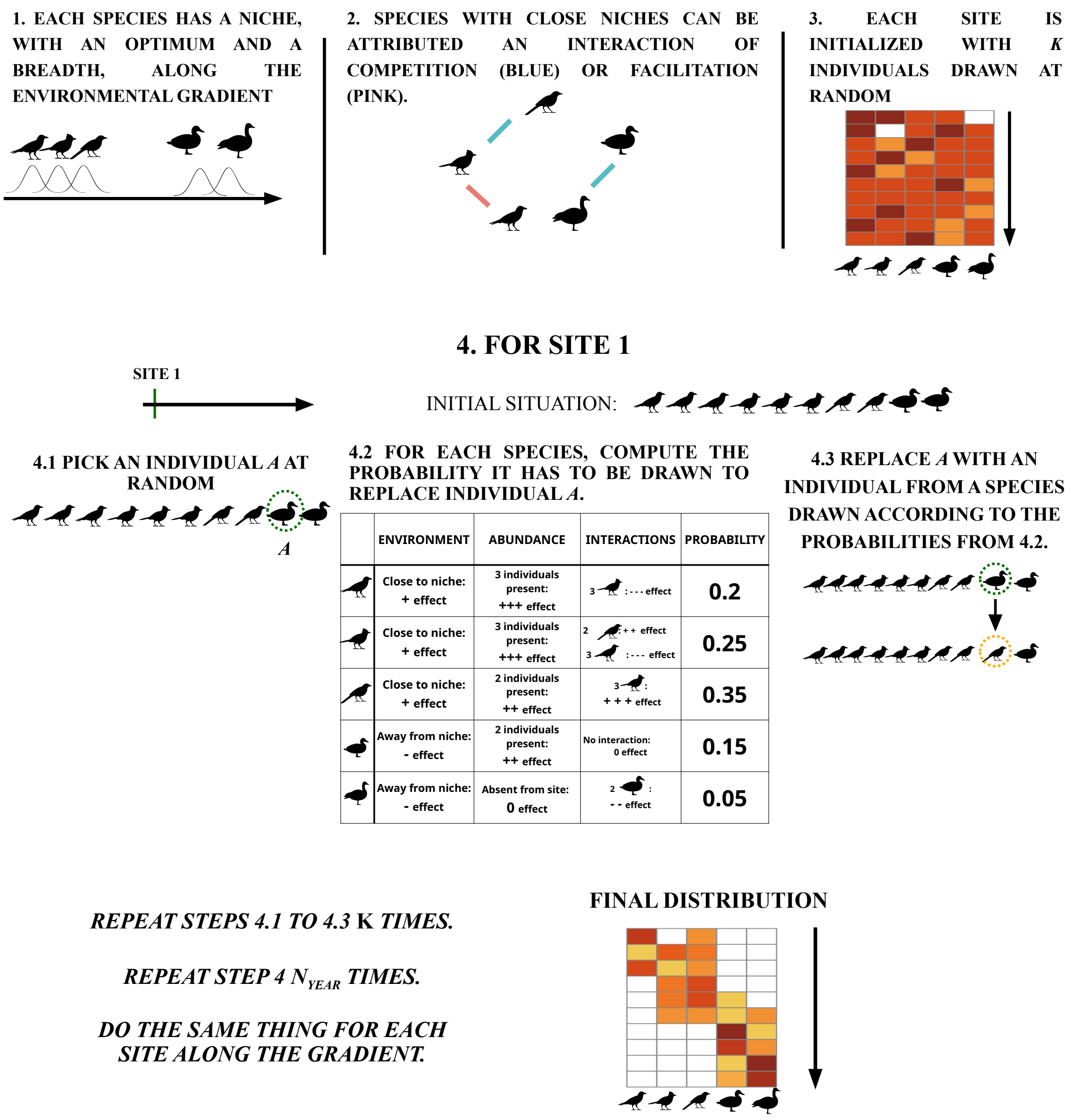


**Figure S1** – Principle of the VirtualCom simulation model, the lottery for space (Section 2.1). Steps 1 and 2, the niches and the interaction network, are shared by the three mechanisms; steps 3 and 4 are the lottery itself.

### S2 The simulated data

Table S1 describes the simulated data. Sites whose counts were proportional to those of another site were removed before fitting; in practice these are sites that hold the same single species.

**Table S1** – The simulated data, by mechanism, niche breadth $b$ and interaction threshold $D$: means over 40 replicates, for the lottery at intensity $a = 2$ and for the Gompertz and Ricker mechanisms at $\kappa = 0.5$ and $\sigma = 0.5$. Sites: number of sites kept, of 50. Share of zeros: in the simulation with interactions (inter.) and in its paired no-interaction control. Individuals per site: mean over the sites, and poorest site. Species per site: number of species present at a site. Prevalence: share of the sites that a species occupies, median over the species. Interactions: number of competition and facilitation pairs in the network. All columns but one describe the simulation with interactions.

| | | | share of zeros | | individuals per site | | species | | interactions | |
|---|---|---|---|---|---|---|---|---|---|---|
| mechanism | $b$ | sites | inter. | control | mean | poorest | per site | prevalence | comp. | facil. |
| *interaction threshold $D = 10$* | | | | | | | | | | |
| Lottery | 2 | 45.8 | 0.92 | 0.92 | 100 | 100 | 1.6 | 0.08 | 6.9 | 3.4 |
| | 5 | 48.5 | 0.87 | 0.86 | 100 | 100 | 2.5 | 0.13 | 7.2 | 3.5 |
| | 10 | 49.6 | 0.79 | 0.75 | 100 | 100 | 4.1 | 0.21 | 7.4 | 3.5 |
| | 20 | 50.0 | 0.64 | 0.57 | 100 | 100 | 7.3 | 0.37 | 6.7 | 3.1 |
| Gompertz | 2 | 43.3 | 0.87 | 0.87 | 122 | 0 | 2.6 | 0.14 | 7.3 | 3.4 |
| | 5 | 48.3 | 0.71 | 0.71 | 287 | 26 | 5.8 | 0.31 | 7.5 | 3.5 |
| | 10 | 49.8 | 0.46 | 0.46 | 538 | 107 | 10.8 | 0.57 | 7.5 | 3.5 |
| | 20 | 50.0 | 0.14 | 0.14 | 993 | 336 | 17.2 | 0.89 | 6.7 | 3.1 |
| Ricker | 2 | 42.8 | 0.89 | 0.87 | 103 | 0 | 2.2 | 0.12 | 7.3 | 3.4 |
| | 5 | 47.9 | 0.74 | 0.71 | 238 | 21 | 5.2 | 0.27 | 7.5 | 3.5 |
| | 10 | 49.9 | 0.50 | 0.47 | 437 | 84 | 9.9 | 0.51 | 7.5 | 3.5 |
| | 20 | 50.0 | 0.17 | 0.15 | 813 | 270 | 16.6 | 0.86 | 6.7 | 3.1 |
| *interaction threshold $D = 50$* | | | | | | | | | | |
| Lottery | 2 | 42.5 | 0.92 | 0.91 | 100 | 100 | 1.6 | 0.08 | 26.3 | 13.8 |
| | 5 | 48.4 | 0.87 | 0.85 | 100 | 100 | 2.6 | 0.13 | 28.6 | 14.3 |
| | 10 | 49.5 | 0.80 | 0.75 | 100 | 100 | 4.0 | 0.20 | 28.7 | 14.2 |
| | 20 | 49.9 | 0.69 | 0.57 | 100 | 100 | 6.2 | 0.31 | 29.2 | 14.4 |
| Gompertz | 2 | 44.0 | 0.87 | 0.87 | 123 | 0 | 2.5 | 0.14 | 28.3 | 13.9 |
| | 5 | 48.1 | 0.70 | 0.70 | 290 | 25 | 5.9 | 0.31 | 29.2 | 14.4 |
| | 10 | 50.0 | 0.47 | 0.47 | 537 | 122 | 10.7 | 0.56 | 29.0 | 14.2 |
| | 20 | 50.0 | 0.14 | 0.14 | 1004 | 291 | 17.3 | 0.89 | 29.4 | 14.5 |
| Ricker | 2 | 43.5 | 0.88 | 0.87 | 103 | 0 | 2.3 | 0.12 | 28.3 | 13.9 |
| | 5 | 47.8 | 0.76 | 0.71 | 237 | 20 | 4.9 | 0.25 | 29.2 | 14.4 |
| | 10 | 50.0 | 0.59 | 0.47 | 426 | 95 | 8.1 | 0.42 | 29.0 | 14.2 |
| | 20 | 50.0 | 0.31 | 0.15 | 768 | 227 | 13.8 | 0.71 | 29.4 | 14.5 |

The niche breadth sets how sparse the data are: the proportion of zero counts goes from about nine in ten with the narrowest niches ($b = 2$), where a site holds about two species, to between 0.14 and two thirds, depending on the mechanism, with the broadest ($b = 20$). The mechanism sets how many individuals are counted: exactly $K = 100$ at every site under the lottery, and from about a hundred to about a thousand per site on average under the Gompertz and Ricker mechanisms, where the sites are unequal and some of them hold no individual at all when the niches are narrow.

Figures S2 and S3 show one replicate: its interaction network, and the counts that the three mechanisms produce from the same species, niches and network, with the interactions and without them. The lottery leaves each species on a narrow range of sites, and its competitive exclusions are visible; under the Gompertz mechanism the two panels can hardly be told apart, the interactions acting on the fluctuations only; under the Ricker mechanism the interactions open gaps in the ranges of some species.

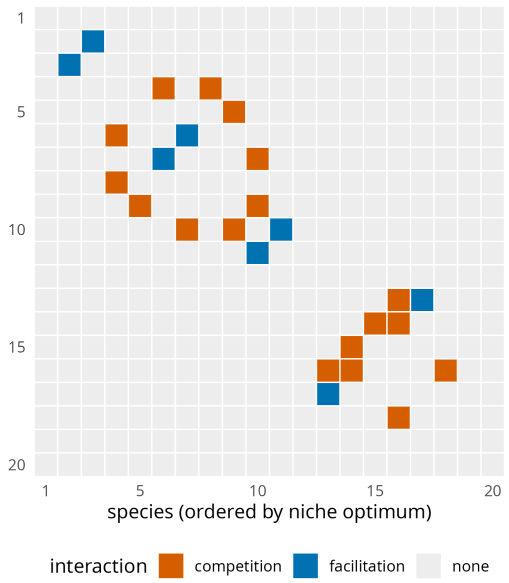


**Figure S2** – The interaction network of one replicate at $b = 10$ and $D = 10$: 9 competition pairs and 4 facilitation pairs among 20 species, ordered by niche optimum. All the interactions lie close to the diagonal: they link species with close optima.

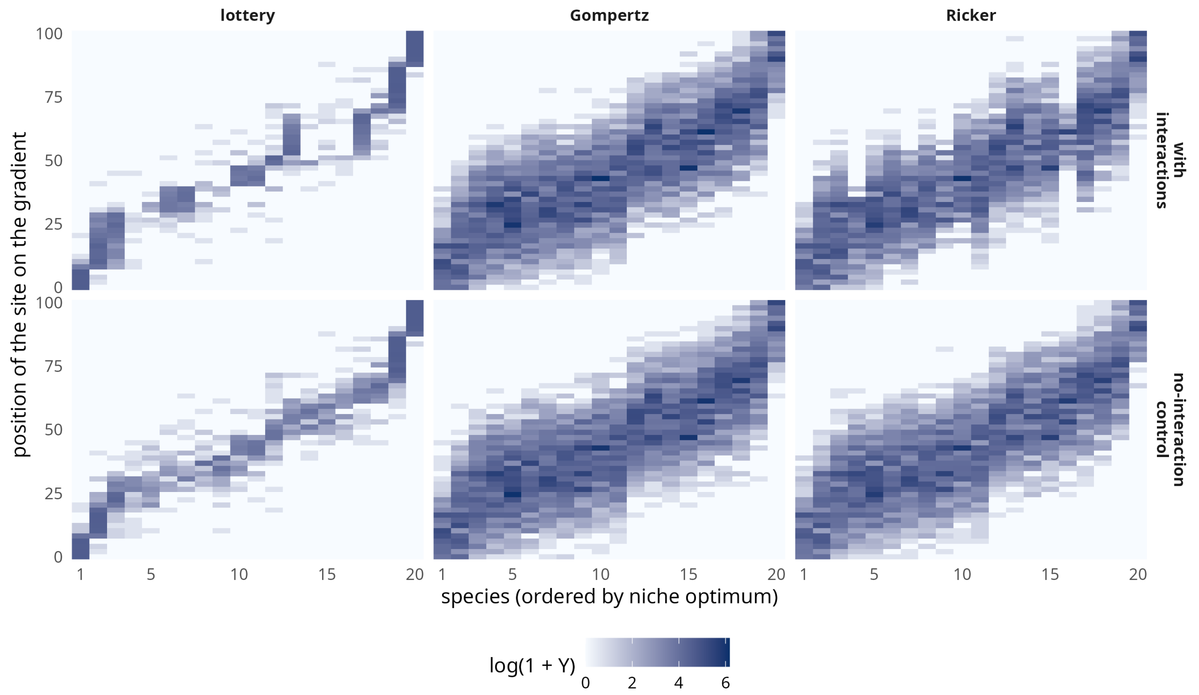


**Figure S3** – The counts of the replicate of Figure S2 under the three mechanisms (lottery at $a = 2$; Gompertz and Ricker at $\kappa = 0.5$ and $\sigma = 0.5$), for the simulation with interactions and its paired no-interaction control. Each cell is a species at a site.

### S3 The models compared

This section gives the mathematical form of the estimators of Section 2.2 and of the two JSDMs kept as points of comparison, with the settings used. Table S2 gathers the notation.

**Table S2** – Notation.

| | |
|---|---|
| $n$, $p$ | numbers of sites and of species |
| $i$; $j$, $k$ | index of a site; indices of species |
| $x_i$ | position of site $i$ on the environmental gradient |
| $Y_{ij}$ | count of species $j$ at site $i$ |
| $b(x) \in \mathbb{R}^d$ | abiotic basis evaluated at $x$ (Table 3), intercept included |
| $\beta_j \in \mathbb{R}^d$, $B = (\beta_1, \dots, \beta_p)$ | abiotic coefficients of species $j$; their $d \times p$ matrix |
| $o_i$ | offset of site $i$ |
| $Z_i \in \mathbb{R}^p$ | Gaussian latent vector of site $i$ |
| $\Sigma$, $\Omega = \Sigma^{-1}$ | covariance and precision matrices of $Z_i$ |
| $q$ | number of latent factors (rank of $\Sigma$) |
| $\lambda$ | penalty of the graphical lasso |
| $W$ | true interaction network of the simulation |

#### A general form

All the models considered here have the same structure. A count is drawn from a distribution whose mean depends, through a logarithmic link, on an abiotic effect and on a Gaussian latent variable:

$$Y_{ij} \mid Z_{ij} \sim \mathcal{F}\left(\exp(o_i + b(x_i)^\top \beta_j + Z_{ij})\right), \qquad Z_i \sim \mathcal{N}(0, \Sigma).$$

The models differ by the distribution $\mathcal{F}$, by the structure imposed on $\Sigma$ or on $\Omega$, and by the method of inference (Table S3). The *latent Gaussian variable* $Z_i$ has one coordinate per species. It must not be confused with the *latent factors* of the models of reduced rank, which are $q < p$ independent Gaussian variables per site of which $Z_i$ is a linear combination.

**Table S3** – The models compared, in the general form of this section.

| Model | Counts | Latent covariance | Inference | Association read |
|---|---|---|---|---|
| log-GGM | Gaussian, on $\log(1 + Y)$ | full rank | least squares | partial |
| PLNPCA | Poisson | $CC^\top$, rank $q$ | variational | partial, after a graphical lasso |
| PLN | Poisson | full rank | variational | partial, after a graphical lasso |
| PLNnetwork | Poisson | sparse precision | variational, penalised | partial |
| gllvm | Poisson | $\Gamma\Gamma^\top$, rank 2 | variational | marginal, or partial after a graphical lasso |
| Hmsc | Poisson, with a Gaussian residual on the log scale | $\Lambda^\top\Lambda$, number of factors adapted | Bayesian, MCMC | marginal |

#### The Poisson log-normal family

*PLN* is model (1), with $\mathcal{F}$ a Poisson distribution and $\Sigma$ unconstrained. The latent layer makes the counts overdispersed and correlated: with $\mu_{ij} = \mathbb{E}(Y_{ij})$,

$$\mathbb{V}(Y_{ij}) = \mu_{ij} + \mu_{ij}^2 \left(e^{\Sigma_{jj}} - 1\right), \qquad \mathrm{Cov}(Y_{ij}, Y_{ik}) = \mu_{ij}\,\mu_{ik}\left(e^{\Sigma_{jk}} - 1\right).$$

The likelihood has no closed form, since $Z_i$ must be integrated out. PLNmodels maximises a variational lower bound instead, in which the conditional distribution of each $Z_i$ given the counts is approximated by a Gaussian distribution with mean $m_i$ and diagonal covariance $\mathrm{diag}(s_i^2)$:

$$J(B, \Sigma, m, s) = \sum_{i=1}^{n} \Big[ Y_i^\top \eta_i - \mathbf{1}^\top \exp\left(\eta_i + s_i^2/2\right) - \tfrac{1}{2}\, m_i^\top \Omega\, m_i - \tfrac{1}{2}\,\mathrm{tr}\left(\Omega\, \mathrm{diag}(s_i^2)\right) + \tfrac{1}{2} \log|\Omega| + \tfrac{1}{2}\, \mathbf{1}^\top \log s_i^2 \Big] + \text{constant}, \qquad \eta_i = o_i \mathbf{1} + B^\top b(x_i) + m_i.$$

*PLNPCA* adds the constraint $\Sigma = CC^\top$, with $C$ a $p \times q$ matrix: $Z_i = C\,U_i$, where $U_i \sim \mathcal{N}(0, I_q)$ are the $q$ latent factors of site $i$. The variational approximation then bears on $U_i$, with mean $m_i \in \mathbb{R}^q$ and covariance $\mathrm{diag}(s_i^2)$. The estimate of $\Sigma$ that PLNPCA returns, and to which the graphical lasso is applied, is the variational one,

$$\hat{\Sigma} = \hat{C} \left( \frac{1}{n} \sum_{i=1}^{n} \left[ m_i m_i^\top + \mathrm{diag}(s_i^2) \right] \right) \hat{C}^\top,$$

which has rank $q$.

*PLNnetwork* maximises the same bound penalised on the precision matrix,

$$J(B, \Omega, m, s) - \lambda \sum_{j \neq k} |\Omega_{jk}|,$$

for a decreasing sequence of penalties $\lambda$. The abiotic coefficients, the variational parameters and the sparse $\Omega$ are thus estimated together.

#### The log-GGM and the two-step estimators

The log-GGM replaces the Poisson layer by a transformation: $\log(1 + Y_{ij}) = b(x_i)^\top \beta_j + \varepsilon_{ij}$, with $\varepsilon_i \sim \mathcal{N}(0, \Sigma)$. The coefficients are estimated by least squares, species by species, and $\Sigma$ by the covariance of the residuals $\hat{\varepsilon}_i$ with the degrees of freedom of the regression, $\hat{\Sigma} = \sum_i \hat{\varepsilon}_i \hat{\varepsilon}_i^\top / (n - d)$.

A two-step estimator then applies the graphical lasso to an estimated covariance $\hat{\Sigma}$, that of the log-GGM, of PLNPCA, of PLN or of gllvm:

$$\hat{\Omega}_\lambda = \arg\max_{\Omega \succ 0}\ \log|\Omega| - \mathrm{tr}(\hat{\Sigma}\,\Omega) - \lambda \sum_{j \neq k} |\Omega_{jk}|.$$

This is the penalised way of inverting a covariance matrix; it applies to a covariance of reduced rank, which has no inverse.

#### gllvm and Hmsc

Both are latent-factor models. In *gllvm*, as we used it,

$$Y_{ij} \mid U_i \sim \mathcal{P}\left(\exp(b(x_i)^\top \beta_j + U_i^\top \gamma_j)\right), \qquad U_i \sim \mathcal{N}(0, I_q),$$

with $q = 2$ latent factors, the default of the package. The covariance of the latent layer is $\Gamma\Gamma^\top$, where $\Gamma$ is the $p \times q$ matrix of the loadings $\gamma_j$: it has rank 2, with no diagonal term added. This is the model of PLNPCA at $q = 2$. The model was fitted by the variational approximation that is the default of the package.

In *Hmsc*, as we used it, the counts are Poisson with a log-normal residual:

$$Y_{ij} \mid L_{ij} \sim \mathcal{P}\left(\exp(L_{ij})\right), \qquad L_{ij} = b(x_i)^\top \beta_j + U_i^\top \lambda_j + \varepsilon_{ij}, \qquad \varepsilon_{ij} \sim \mathcal{N}(0, \sigma_j^2),$$

with latent factors $U_i$ at the level of the site. Three features set it apart. The abiotic coefficients share information between species: they have a common Gaussian prior, $\beta_j \sim \mathcal{N}(\gamma, V)$, whose mean and covariance are estimated. The number of factors is not fixed: the loadings have a shrinkage prior, and factors are added or removed during the burn-in. And the inference is Bayesian, by Markov chain Monte Carlo. The network is read in the matrix of associations $\Lambda^\top\Lambda$, turned into correlations: its posterior mean, and for each pair the posterior probability that the association is positive. A network is obtained by keeping the pairs for which this probability is above 0.9 or below 0.1; this is the only form of sparsity, and it comes after the fit.

#### Settings

Table S4 lists the settings. Whatever the basis, a model was fitted with the same settings. A PLN fit that failed was tried again with the second optimiser of the package, then without profiling the covariance out of the objective; if it failed again, PLNnetwork was started from a PLN fit with a diagonal covariance, and levels 3 and the marginal reading were recorded as missing. A fit was stopped after ten minutes.

Fits failed with one basis only, the one that ignores the environment: between 0.7% (lottery) and 4.2% (Ricker) of the fits of PLN and of PLNnetwork, presumably because the latent layer then has to carry the whole response to the gradient, which leaves the covariance of the PLN model poorly conditioned. No fit of PLNPCA failed, and four fits of gllvm did, of 48 000.

**Table S4** – Settings of the models. All the computations were run with R 4.6.1.

| Model | Function (package) | Settings |
|---|---|---|
| PLN | `PLN` (PLNmodels 1.3.2) | defaults |
| PLNPCA | `PLNPCA` (PLNmodels) | ranks 2, 5 and 10; defaults otherwise |
| PLNnetwork | `PLNnetwork` (PLNmodels) | started from the PLN fit; 76 penalties log-spaced over three decades; diagonal of $\Omega$ not penalised |
| Graphical lasso | `graphical_lasso` (PLNmodels) | 101 penalties log-spaced over four decades, from $\max_{j \neq k} \lvert\hat{\Sigma}_{jk}\rvert$ |
| log-GGM | `lm.fit` (stats), then the graphical lasso | — |
| gllvm | `gllvm` (gllvm 2.0.13) | Poisson family; 2 latent variables; variational approximation; defaults otherwise |
| Hmsc | `Hmsc`, `sampleMcmc` (Hmsc 3.3.7) | log-normal Poisson; one random level at the site; 1000 samples kept every 10 iterations after a burn-in of 5000 |

### S4 Checks on the simulators and on the scoring

#### Convergence of the lottery

The lottery starts from a community drawn at random and is run for a number of "years", each year replacing as many individuals as a site holds. We ran it for 10 years. To check that this is enough, we ran it for 40 years in the eight configurations and recorded, year after year, the change in the abundance of a species at a site (Figure S4). The change is large in the first years, 3.5 to 5.8 individuals per species and site in the first one, while the random initial community is sorted by the niches. It then settles on a plateau, which is not zero since the lottery keeps replacing individuals: 0.14 to 1.05 at year 10, depending on the niche breadth, against 0.10 to 0.88 at year 40. Ten years thus bring the communities close to their stationary regime in every configuration, with a small residual drift when the niches are narrow; the simulation and its control behave alike. The Gompertz and Ricker dynamics were run for 200 steps. In the Ricker dynamics, the exponent of the growth factor was capped at 3 for numerical safety; the cap only acts in degenerate cases, when a facilitator is abundant at the edge of the niche of the species it helps.

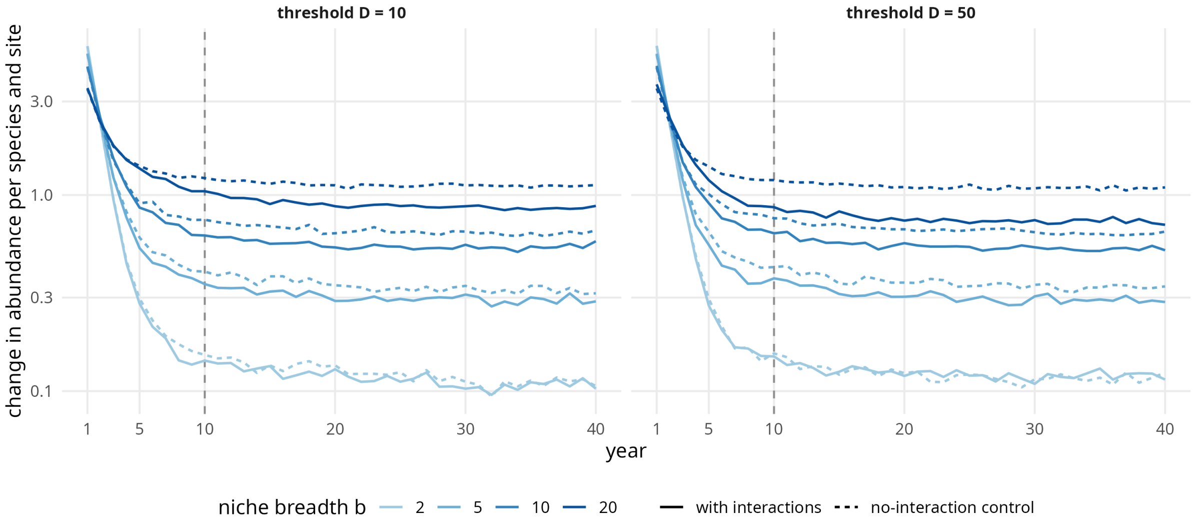


**Figure S4** – Convergence of the lottery. Absolute change in the abundance of a species at a site from one year to the next, averaged over the species, the sites and 10 replicates, in the eight configurations (lottery at $a = 2$; logarithmic scale). Dashed line: the 10 years used in the paper.

#### The relative abundance index

Before considering JSDMs, one can first wonder whether the patterns in the simulated data correspond to what one would intuitively expect given the way VirtualCom was designed: given two species with close niche optima, they are expected to be found together less (resp. more) than the other pairs species with similar niche proximity if there is a competition (resp. facilitation) interaction between them. To test this correspondence Si-moussi et al. (2020) proposed a Relative Abundance Index (RAI). For two species $j$, $k$, when $n$ sites are considered along the environmental gradient, we define:

$$\begin{aligned} \mathrm{RAI}_{jk} &= \mathrm{avg}\left(\{\delta_{i,j} | 1 \leq i \leq n \, y_{ij} > 0, y_{ik} > 0\}\right) \\ \text{with } \delta_{i,j} &= y_{ij} - \bar{y}_j, \end{aligned}$$

with $y_{ij}$ the abundance of species $j$ at site $i$ and $\bar{y}_j$ its average abundance over all the sites. This asymmetric index is the empirical transcription of the idea that *if $j$ is in competition with $k$, then when both species are present $j$'s abundance should be lower than it is on average*. However, it fails to account for the situation in which the competition is so strong that the two species are never found together. It is essentially an empirical test to ensure that the simulation "does what is says" and is correctly parameterized.

The RAIs are measured separately for four categories of species pairs: competing species, facilitating species, species with no interactions but close niches (below $D_{\mathrm{threshold}}$ so that they should still often occur together) and species with no interactions and distant niches.

Table S5 gives the median of this index by type of pair. Pairs of species with close niches and no interaction have a large positive index under every mechanism: both species are present, and above their average abundance, at the sites that suit them both. The index is computed on raw abundances and mostly measures this shared response to the environment, which is also why it is positive for pairs in competition. An index near zero when one species is present almost everywhere, and undefined when the two never meet, are its other limits. Besides, the index clearly separates competition from facilitation, on either side of the pairs with no interaction, under the lottery only. Under the Ricker mechanism facilitation stands out with narrow niches and nothing else does. Under the Gompertz mechanism the three types of pairs cannot be told apart: the interactions shape the fluctuations around the level set by the environment, and not that level. A summary of the raw data can therefore be blind to interactions that an estimator of the dependence between species captures.

**Table S5** – Relative abundance index by type of pair: median over the pairs and the replicates, simulation with interactions, interaction threshold $D = 10$. Pairs with no interaction are split into those whose niche optima are closer than the threshold and the others.

| | | | | no interaction | |
|---|---|---|---|---|---|
| mechanism | $b$ | competition | facilitation | close niches | distant niches |
| Lottery | 2 | 0.9 | 41.3 | 29.6 | 15.4 |
| | 5 | 1.5 | 32.2 | 22.1 | 0.7 |
| | 10 | 1.8 | 24.6 | 13.3 | 3.1 |
| | 20 | 1.9 | 15.0 | 6.7 | 2.8 |
| Gompertz | 2 | 37.8 | 39.8 | 37.6 | -2.5 |
| | 5 | 40.9 | 39.2 | 40.6 | 11.7 |
| | 10 | 25.0 | 24.9 | 24.7 | 20.2 |
| | 20 | 7.2 | 7.0 | 7.2 | 7.5 |
| Ricker | 2 | 30.4 | 48.4 | 35.0 | -1.7 |
| | 5 | 40.5 | 49.8 | 39.1 | 10.9 |
| | 10 | 26.8 | 28.6 | 24.5 | 16.6 |
| | 20 | 9.1 | 7.4 | 7.5 | 6.9 |

#### Reference networks

Table S6 gives the raw AUC of three rankings that use no count data. A random network, with or without the true numbers of interactions, is at the chance level. Ranking the pairs by the proximity of their niche optima is far above it, because the interactions are drawn among the pairs with close niches: this is the part of a raw score that owes nothing to the interactions, and the reason for reading increments over the paired control.

**Table S6** – Raw AUC of reference rankings, mean over the mechanisms and the replicates of the main design. Niche proximity: the pairs are ranked by the proximity of their niche optima.

| | $D = 10$ | | $D = 50$ | |
|---|---|---|---|---|
| reference | competition | facilitation | competition | facilitation |
| Random network | 0.50 | 0.50 | 0.50 | 0.50 |
| Random network of the true density | 0.51 | 0.50 | 0.49 | 0.48 |
| Niche proximity | 0.91 | 0.90 | 0.63 | 0.62 |

### S5 Supplementary experiments

Each experiment relaxes one assumption of the main design (Table S7), under the lottery at $a = 2$ and under the Gompertz mechanism at $\kappa = 0.5$ and $\sigma = 0.5$, with $b = 10$ and $D = 10$. The tables give the increments of level 1 (log-GGM) and level 4 (PLNnetwork) for competition, each at its best basis; the figures add facilitation and the intervals.

**Table S7** – The supplementary experiments, with $n = 50$ sites and $p = 20$ species unless it is the factor varied.

| Experiment | Settings | Factor varied | Estimators |
|---|---|---|---|
| Imperfect covariate | lottery and Gompertz; $b = 10$, $D = 10$ | error on the covariate, 0 to 1 niche breadth; with or without a second, unmeasured gradient | levels 1 and 4 |
| Sites and species | lottery and Gompertz; $b = 10$, $D = 10$ | $n \in \{30, 50, 100\}$, $p \in \{20, 40\}$ | levels 1 and 4 |
| Dense network | lottery and Gompertz; $b = 10$, $D = 10$ | network of rank 1, 3 or 10 | levels 1 and 4, gllvm |

#### An imperfect covariate, an unmeasured gradient

The estimators were given the gradient with a Gaussian error, of standard deviation 0 to 1 niche breadth. Independently, a second gradient could act on the communities without being given to the estimators: each species has a second niche, of the same breadth, on a gradient that is independent of the first.

Under the Gompertz mechanism, the advantage of PLNnetwork requires a nearly perfect covariate (Table S8, Figure S5). It holds with no error and with an error of 0.1 niche breadth, and has vanished at 0.25 for the AUC; the top-$k$ precision of both estimators falls to zero from 0.5 on. Under the lottery the AUC resists an error of half a niche breadth, and the top-$k$ precision an error of a quarter. A second, unmeasured gradient removes most of what there was to capture under both mechanisms and for both estimators.

In raw scores (Figure S6), under the Gompertz mechanism, from an error of 0.25 niche breadth on, the raw AUC of competition falls below 0.5 with interactions and in the control alike: what is left of the niches in the residual associations ranks the pairs with close niches the wrong way round. The increment can then remain positive while the ranking is worse than a random one.

**Table S8** – Imperfect covariate and unmeasured gradient: increments for competition. Error: standard deviation of the error on the covariate, in niche breadths.

| | | | AUC | | top-$k$ precision | |
|---|---|---|---|---|---|---|
| mechanism | second gradient | error | log-GGM | PLNnetwork | log-GGM | PLNnetwork |
| Lottery | no | 0 | 0.321 | 0.328 | 0.278 | 0.309 |
| | no | 0.1 | 0.325 | 0.320 | 0.304 | 0.319 |
| | no | 0.25 | 0.339 | 0.329 | 0.288 | 0.334 |
| | no | 0.5 | 0.365 | 0.374 | 0.178 | 0.233 |
| | no | 1 | 0.376 | 0.297 | 0.071 | 0.062 |
| | yes | 0 | 0.112 | 0.079 | 0.016 | 0.017 |
| | yes | 0.1 | 0.112 | 0.073 | 0.024 | 0.042 |
| | yes | 0.25 | 0.112 | 0.082 | 0.018 | 0.031 |
| | yes | 0.5 | 0.115 | 0.087 | 0.016 | 0.008 |
| | yes | 1 | 0.114 | 0.096 | 0.016 | 0.012 |
| Gompertz | no | 0 | 0.137 | 0.173 | 0.115 | 0.180 |
| | no | 0.1 | 0.121 | 0.150 | 0.092 | 0.109 |
| | no | 0.25 | 0.104 | 0.103 | 0.022 | 0.037 |
| | no | 0.5 | 0.063 | 0.067 | 0.000 | 0.000 |
| | no | 1 | 0.051 | 0.051 | 0.000 | 0.000 |
| | yes | 0 | 0.020 | 0.007 | 0.022 | 0.003 |
| | yes | 0.1 | 0.017 | 0.004 | 0.015 | 0.001 |
| | yes | 0.25 | 0.019 | 0.010 | 0.009 | 0.013 |
| | yes | 0.5 | 0.008 | 0.012 | 0.007 | 0.011 |
| | yes | 1 | 0.017 | -0.002 | 0.016 | 0.022 |

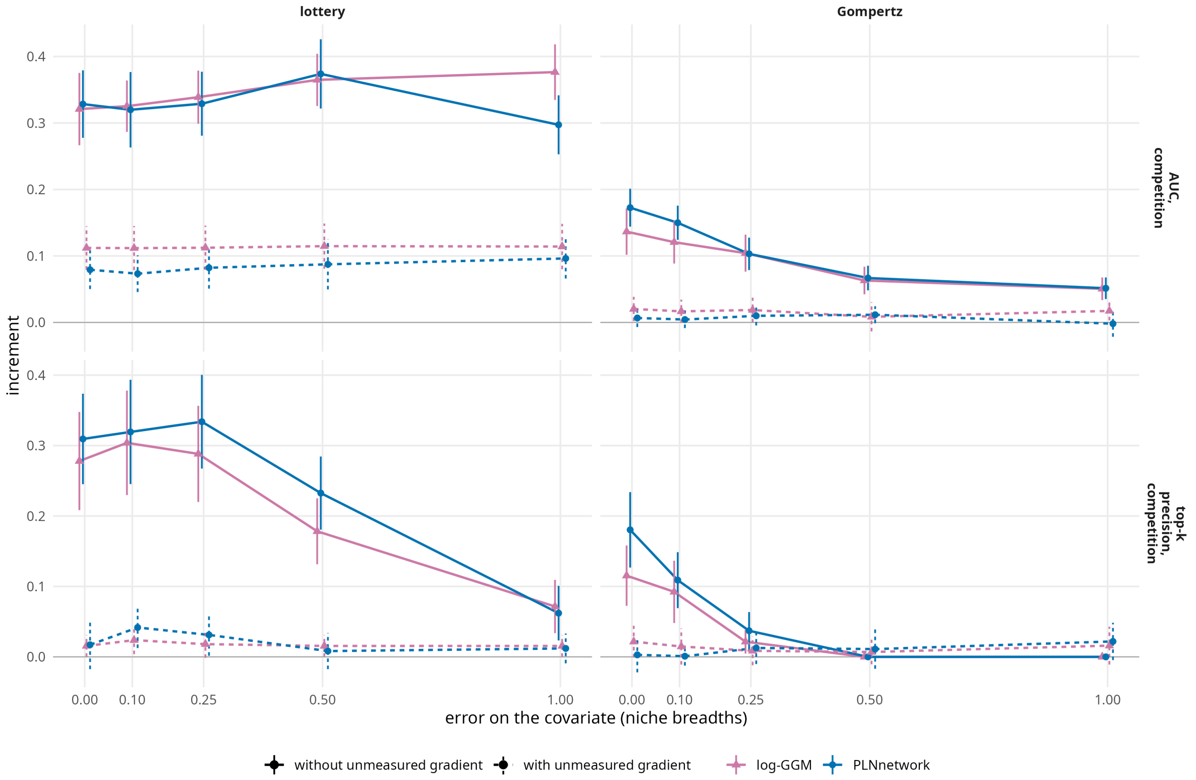


**Figure S5** – Increment against the error on the covariate, without and with a second, unmeasured gradient. Bars: 95% confidence intervals over the 40 replicates.

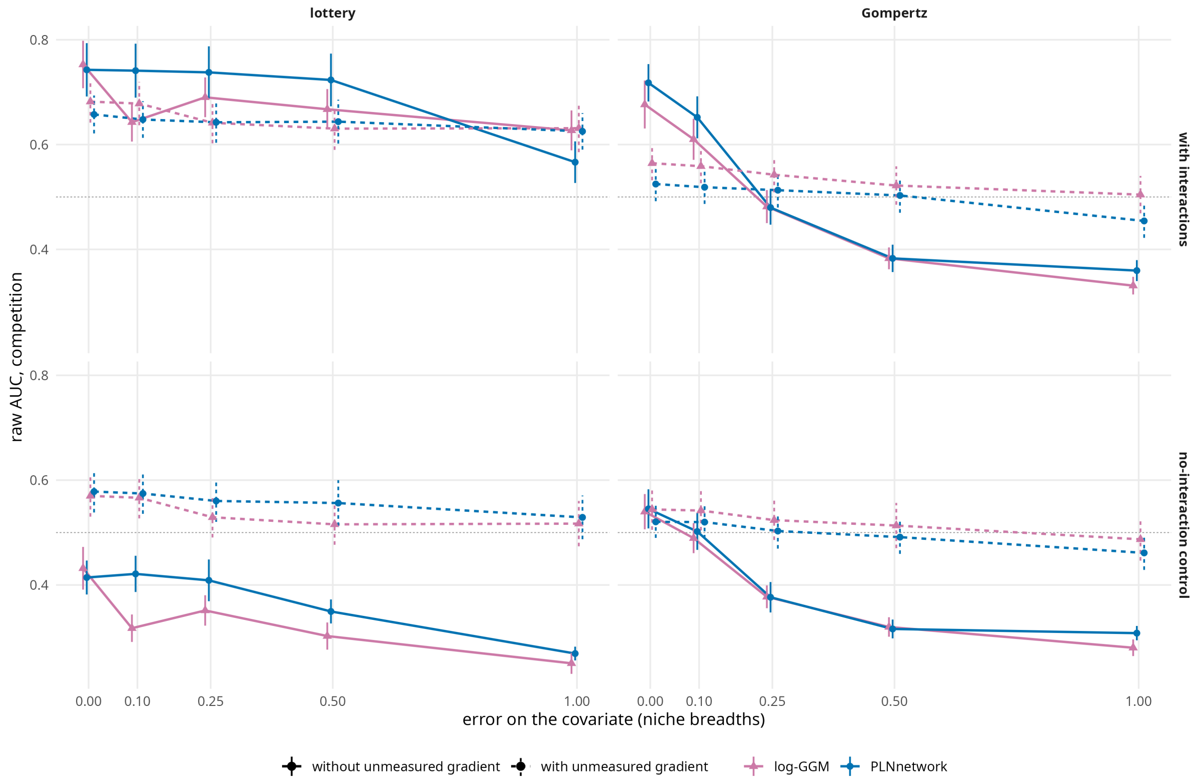


**Figure S6** – The experiment of Figure S5 in raw AUC, for the simulation with interactions and for its control.

#### Numbers of sites and of species

With $n \in \{30, 50, 100\}$ sites and $p \in \{20, 40\}$ species, the increments of both estimators grow with the number of sites and shrink with the number of species (Table S9, Figure S7). Under the Gompertz mechanism PLNnetwork is ahead with 20 species and hardly so with 40. Under the lottery the log-GGM has the better AUC, most clearly with few sites, and PLNnetwork the better top-$k$ precision with 20 species and 50 or 100 sites.

**Table S9** – Numbers of sites $n$ and of species $p$: increments for competition.

| | | | AUC | | top-$k$ precision | |
|---|---|---|---|---|---|---|
| mechanism | $p$ | $n$ | log-GGM | PLNnetwork | log-GGM | PLNnetwork |
| Lottery | 20 | 30 | 0.308 | 0.256 | 0.272 | 0.263 |
| | 20 | 50 | 0.321 | 0.328 | 0.278 | 0.309 |
| | 20 | 100 | 0.479 | 0.464 | 0.398 | 0.478 |
| | 40 | 30 | 0.236 | 0.147 | 0.148 | 0.130 |
| | 40 | 50 | 0.269 | 0.235 | 0.194 | 0.198 |
| | 40 | 100 | 0.343 | 0.332 | 0.275 | 0.280 |
| Gompertz | 20 | 30 | 0.085 | 0.121 | 0.077 | 0.084 |
| | 20 | 50 | 0.137 | 0.173 | 0.115 | 0.180 |
| | 20 | 100 | 0.235 | 0.263 | 0.222 | 0.260 |
| | 40 | 30 | 0.068 | 0.068 | 0.042 | 0.056 |
| | 40 | 50 | 0.096 | 0.101 | 0.083 | 0.092 |
| | 40 | 100 | 0.146 | 0.160 | 0.109 | 0.121 |

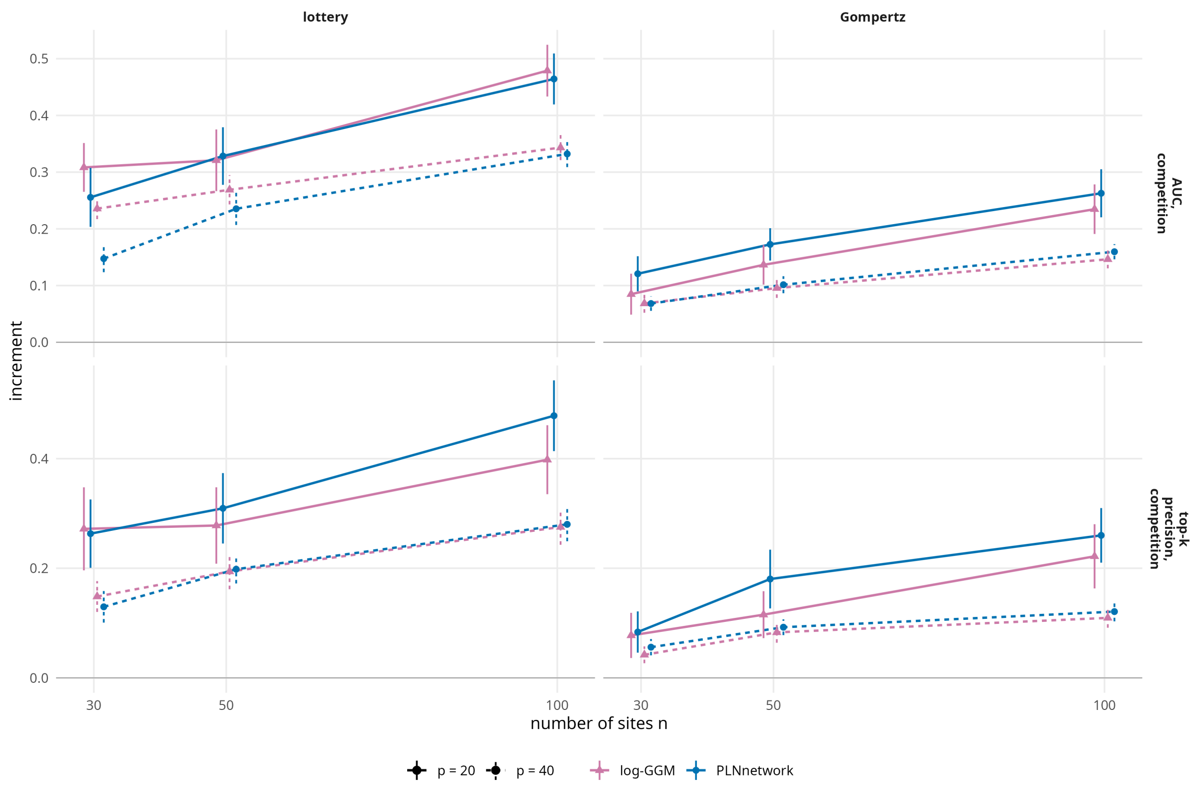


**Figure S7** – Increment against the number of sites, for 20 and 40 species.

#### A dense network of low rank

A sparse network suits an estimator that looks for a sparse precision matrix. Here the network that acts on the communities is dense and of rank 1, 3 or 10: it is built from the leading eigenvectors of the matrix of niche proximities, the first one excepted, which gives interactions of both signs between all the pairs. The estimators were scored on its strongest entries, as many as there are interactions in the sparse design. This is harder for all the estimators under the Gompertz mechanism (Table S10, Figure S8). The log-GGM is slightly ahead for the AUC and PLNnetwork for the top-$k$ precision, as in the main design. gllvm, whose latent factors parameterise exactly a structure of low rank, stays behind both wherever there is something to capture, by a factor of two for the AUC under the lottery: a dense network of low rank does not turn the comparison against the estimator of a sparse precision matrix, nor in favour of a latent-factor model with two factors.

**Table S10** – Dense network of low rank: increments for competition. gllvm: two latent variables, read through the graphical lasso.

| | | AUC | | | top-$k$ precision | | |
|---|---|---|---|---|---|---|---|
| mechanism | rank | gllvm | log-GGM | PLNnetwork | gllvm | log-GGM | PLNnetwork |
| Lottery | 1 | 0.154 | 0.298 | 0.289 | 0.213 | 0.224 | 0.286 |
| | 3 | 0.157 | 0.421 | 0.386 | 0.161 | 0.313 | 0.353 |
| | 10 | 0.218 | 0.488 | 0.449 | 0.149 | 0.354 | 0.376 |
| Gompertz | 1 | 0.012 | 0.063 | 0.051 | 0.018 | 0.013 | 0.014 |
| | 3 | 0.038 | 0.058 | 0.049 | 0.026 | 0.022 | 0.054 |
| | 10 | 0.028 | 0.083 | 0.078 | 0.015 | 0.057 | 0.077 |

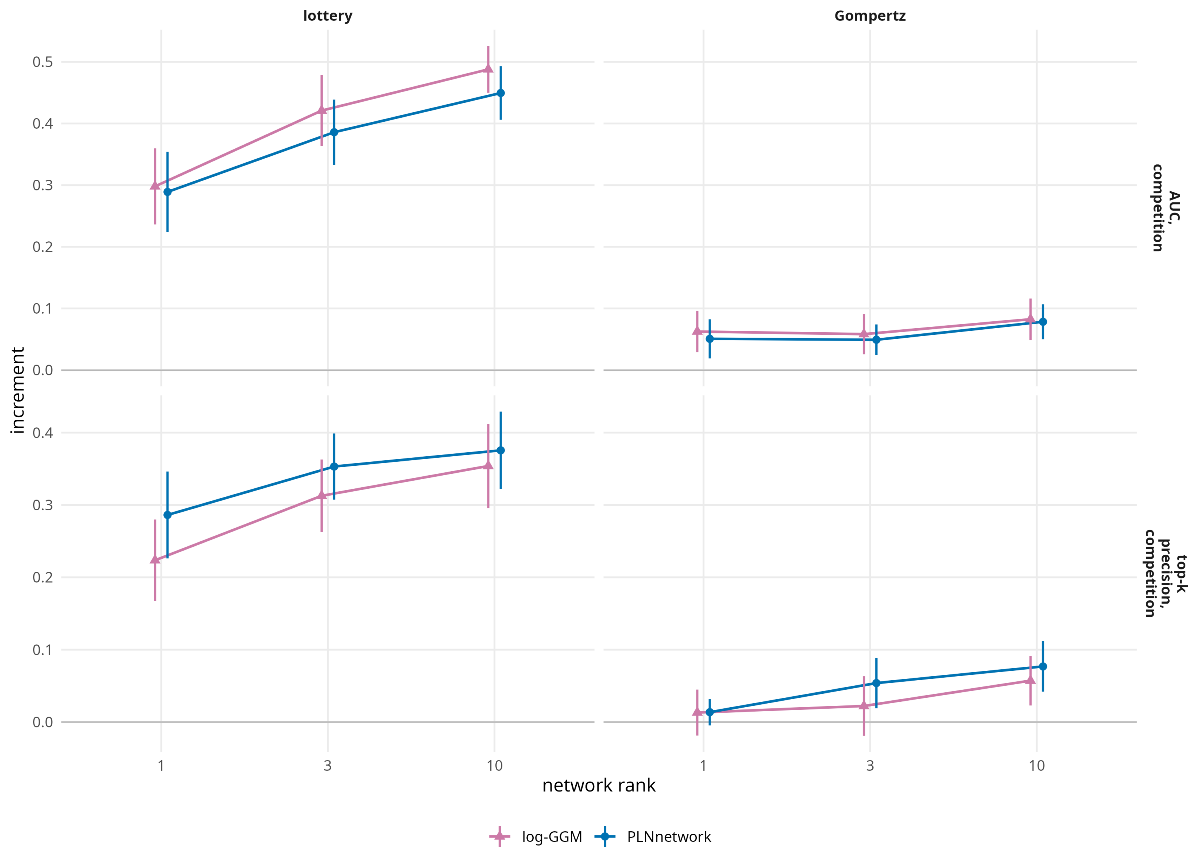


**Figure S8** – Increment against the rank of the true network.

### S6 gllvm and Hmsc

#### gllvm

gllvm with its two latent variables was fitted in the main design and read like the two-step estimators. It is the same model as PLNPCA at rank 2 (Section S3), and the two give the same results (Table S11): the differences are within 0.01 and their signs vary with the mechanism and the metric. A latent-factor JSDM is thus represented in the main text by PLNPCA, which is fitted several times faster and whose rank can be raised at little cost (Section S8).

**Table S11** – PLNPCA at rank 2 minus gllvm with 2 latent variables, both followed by the graphical lasso: difference of increments, mean over the settings of a mechanism and half-width of its 95% confidence interval.

| metric | Lottery | Gompertz | Ricker |
|---|---|---|---|
| AUC, competition | $+0.011 \pm 0.009$ | $-0.001 \pm 0.004$ | $-0.002 \pm 0.005$ |
| AUC, facilitation | $+0.010 \pm 0.009$ | $-0.004 \pm 0.005$ | $-0.002 \pm 0.004$ |
| top-$k$ precision, competition | $+0.004 \pm 0.007$ | $-0.003 \pm 0.005$ | $+0.004 \pm 0.005$ |
| top-$k$ precision, facilitation | $-0.010 \pm 0.009$ | $-0.009 \pm 0.006$ | $-0.005 \pm 0.006$ |

#### gllvm with a negative binomial distribution

To see whether the emission distribution of the JSDM is what limits it under the lottery and the Ricker mechanism, gllvm was fitted with a Poisson and with a negative binomial distribution, at the same number of latent variables (five, and ten for the negative binomial), and read through the same graphical lasso, in nine settings: the three mechanisms (lottery at $a = 2$, Gompertz and Ricker at $\kappa = 0.5$ and $\sigma = 0.5$), $b \in \{5, 10, 20\}$ and $D = 10$, with three bases (categorical, cubic polynomial, spline) and 40 replicates.

Under the Gompertz and the Ricker mechanisms, the two distributions give similar increments, the negative binomial being slightly ahead for the top-$k$ precision under the Gompertz mechanism (Table S12). Under the lottery, the negative binomial does much worse. gllvm with a Poisson distribution is close to PLNPCA at the same rank, and the log-GGM is ahead of all of them under the lottery and the Ricker mechanism. A change of emission distribution therefore does not explain the results of the main text.

**Table S12** – The emission distribution of a latent-factor JSDM: increments for competition, each estimator at its best basis (categorical, cubic polynomial or spline), mean over $b \in \{5, 10, 20\}$ at $D = 10$ (lottery at $a = 2$; Gompertz and Ricker at $\kappa = 0.5$, $\sigma = 0.5$; 40 replicates). In brackets, the number of latent variables (the rank); P: Poisson; NB: negative binomial. gllvm and PLNPCA are read through the graphical lasso.

| metric | mechanism | log-GGM | PLNPCA (5) | gllvm P (5) | gllvm NB (5) | gllvm NB (10) |
|---|---|---|---|---|---|---|
| AUC | Lottery | 0.29 | 0.25 | 0.24 | 0.09 | 0.06 |
| | Gompertz | 0.16 | 0.12 | 0.12 | 0.12 | 0.11 |
| | Ricker | 0.18 | 0.13 | 0.12 | 0.13 | 0.12 |
| top-$k$ precision | Lottery | 0.23 | 0.26 | 0.22 | 0.06 | 0.03 |
| | Gompertz | 0.14 | 0.05 | 0.04 | 0.08 | 0.09 |
| | Ricker | 0.16 | 0.09 | 0.10 | 0.08 | 0.09 |

#### Hmsc

Hmsc is not part of the comparison, because its sampler did not converge in our hands. A single chain per data set leaves no way of seeing it. We ran four chains on each of 30 data sets simulated under the lottery with the narrowest niches ($b = 2$, $D = 10$), with the settings of Table S4, and again with chains five times longer (Table S13). The potential scale reduction factor exceeded 1.05 for more than half of the associations and for most of the abiotic coefficients, and for 23 data sets of 30 its median over the associations was above 1.05. Longer chains did not help: 27 data sets of 30 remained above 1.05.

The consequence for the scores is direct: the standard deviation of the AUC of competition was 0.09 between the chains of a data set and 0.02 between data sets, so that 95% of the variance of the AUC of Hmsc with a single chain comes from the sampler. In a first check on three data sets, the effective sample size of the associations was high within each chain, about 3000 of 4000 draws: each chain mixes well around its own mode, which is why a single chain looks healthy. We did not find settings that make Hmsc converge on these data, and we do not conclude anything about its ability to capture interactions.

**Table S13** – Convergence of Hmsc: four chains on each of 30 data sets simulated under the lottery (niche breadth 2, interaction threshold 10), with the settings of Table S4 and with chains five times longer. Scale reduction factor: potential scale reduction factor over the four chains. The AUC is the one of our first analysis, computed before the corrections described in Section S7; s.d.: standard deviation. Its standard deviation between data sets is the one that remains once the noise of the chains is removed.

| | settings of Table S4 | chains five times longer |
|---|---|---|
| Scale reduction factor, associations: median | 1.08 | 1.14 |
| share above 1.05 | 56% | 67% |
| Scale reduction factor, abiotic coefficients: median | 1.22 | 1.11 |
| share above 1.05 | 93% | 76% |
| Data sets not converged, of 30 | 23 | 27 |
| AUC of competition: s.d. between chains | 0.09 | 0.10 |
| AUC of competition: s.d. between data sets | 0.02 | 0.01 |
| Share of the variance of the AUC due to the sampler | 95% | 99% |

### S7 Pitfalls in scoring an estimated network

We met each of the following pitfalls when scoring an estimated network against a simulated one. Each of them moves the scores by as much as the differences between estimators that one wants to measure. We list them as advice.

*A raw score rewards the environment.* When interactions are drawn among species with close niches, which is the realistic case, a ranking by niche proximity already has a high AUC (Table S6). Score the estimators against a paired control without interactions.

*An AUC read along a path depends on the path.* A path that stops too early leaves true interactions out, tied at the bottom of the ranking: with two decades of penalties, the densest network of PLNnetwork held about two thirds of the pairs under the lottery with the narrowest niches. A curve closed before a recall of one, a path along which an edge can leave after entering, and tied pairs all change the area. Compute the AUC from an explicit ranking of the pairs, with a rule for ties, and give every estimator the same grid.

*Precision, recall and F1 at one threshold compare selection rules.* A network selected by a criterion reflects that criterion as much as the estimator. Compare rankings, or networks of the same density.

### S8 Run times

Table S14 gives the time of one fit, that is one basis on one data set with $n = 50$ sites and $p = 20$ species. The log-GGM is fitted in a few hundredths of a second, the count models in 0.6 to 7 seconds. The time of the graphical lasso of the two-step estimators is not included. The fits were run on a computing server with two Intel Xeon E5-2630 v3 processors (2.40 GHz, 8 cores each), on one thread each.

### S9 Additional results of the main design

This section gathers the figures and tables that the Results of the main text refer to.

**Table S14** – Run time of one fit, in seconds: median and 90% quantile over the fits of the main design.

| | Lottery | | Gompertz | | Ricker | |
|---|---|---|---|---|---|---|
| estimator | median | 90% | median | 90% | median | 90% |
| log-GGM (level 1) | 0.03 | 0.04 | 0.04 | 0.05 | 0.04 | 0.05 |
| PLNPCA, rank 2 | 0.72 | 1.03 | 0.64 | 0.97 | 0.66 | 0.98 |
| PLNPCA, rank 5 | 1.19 | 1.61 | 0.95 | 1.58 | 0.99 | 1.59 |
| PLNPCA, rank 10 (level 2) | 2.00 | 2.50 | 1.42 | 2.54 | 1.51 | 2.55 |
| PLN, then PLNnetwork (levels 3, 4) | 6.70 | 12.13 | 4.15 | 12.65 | 4.59 | 13.09 |
| gllvm, 2 latent variables | 5.80 | 12.76 | 4.39 | 10.00 | 4.74 | 10.70 |

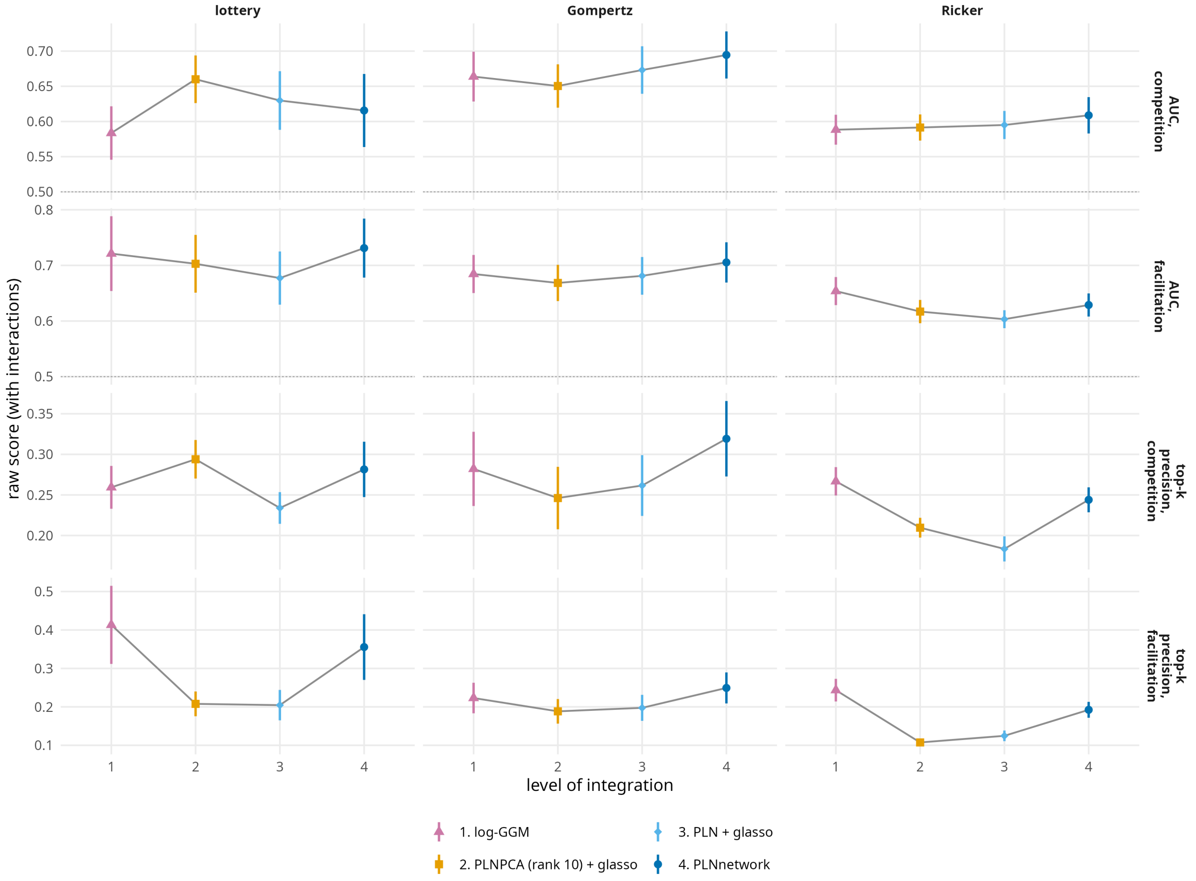


**Figure S9** – The four levels of integration in raw scores, on the simulations with interactions (Figure 2 gives the increments). Grey line: 0.5, the chance level of the AUC.

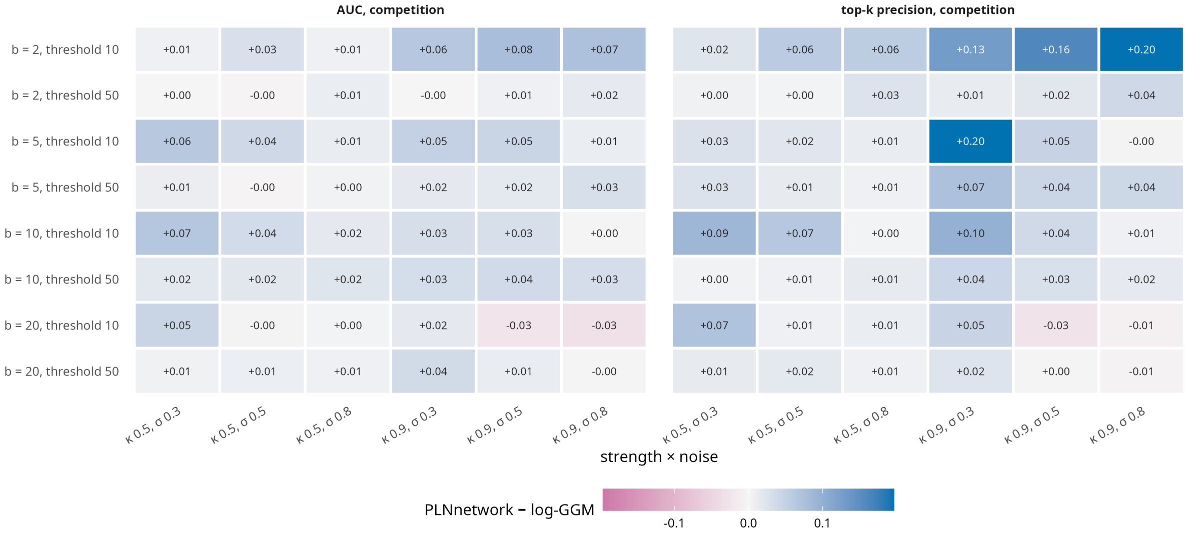


**Figure S10** – Advantage of PLNnetwork over the log-GGM under the Gompertz mechanism: increment of PLNnetwork minus increment of the log-GGM, by configuration and by strength of the interactions ($\kappa$) and process noise ($\sigma$).

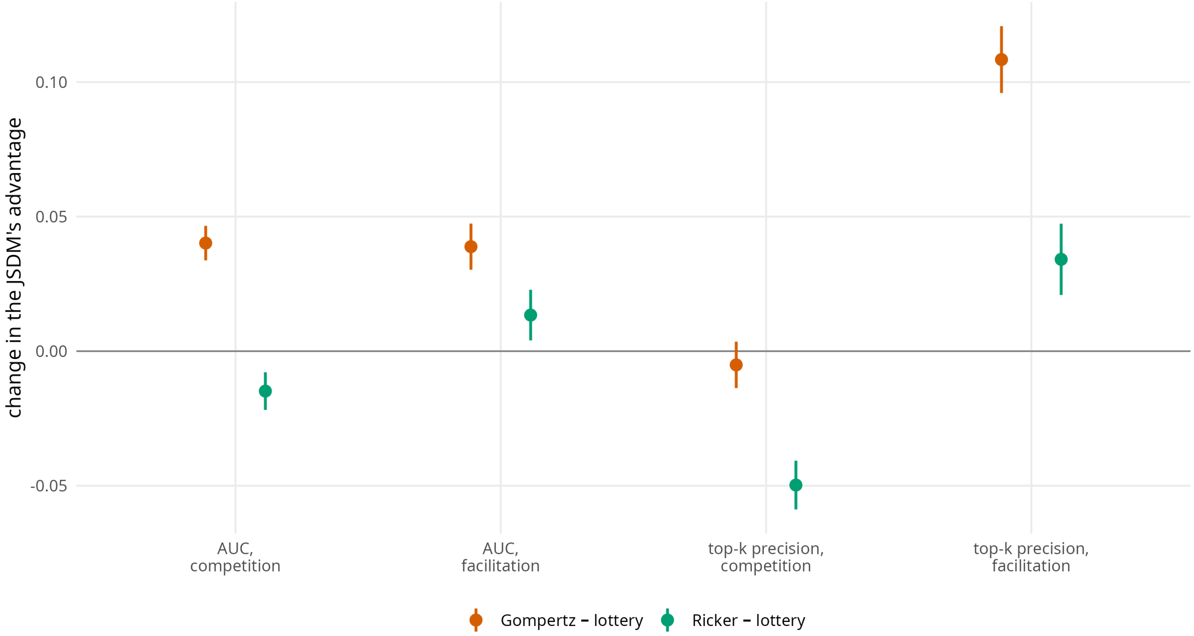


**Figure S11** – The paired effect of the mechanism. Advantage of PLNnetwork over the log-GGM under the Gompertz or the Ricker mechanism, minus the same advantage under the lottery at $a = 2$, on the same configuration, network and replicate; each estimator at its best basis. Bars: 95% confidence intervals over the pairs of replicates.

**Table S15** – Abundance under the Gompertz mechanism: increments for competition of levels 1, 3 and 4, each at its best basis among four (categorical, quadratic, cubic polynomial, spline), against the constant $K$ that multiplies the expected counts. $b = 10$, $D = 10$, $\sigma = 0.5$; the same communities at every $K$, and $K = 100$ is the main design.

| | | AUC | | | top-$k$ precision | | |
|---|---|---|---|---|---|---|---|
| $\kappa$ | $K$ | log-GGM | PLN + glasso | PLNnetwork | log-GGM | PLN + glasso | PLNnetwork |
| 0.5 | 3 | 0.053 | 0.061 | 0.058 | 0.024 | 0.044 | 0.049 |
| 0.5 | 10 | 0.099 | 0.139 | 0.150 | 0.094 | 0.077 | 0.122 |
| 0.5 | 30 | 0.153 | 0.171 | 0.193 | 0.135 | 0.074 | 0.169 |
| 0.5 | 100 | 0.137 | 0.149 | 0.173 | 0.115 | 0.065 | 0.180 |
| 0.5 | 1000 | 0.171 | 0.148 | 0.199 | 0.159 | 0.049 | 0.209 |
| 0.9 | 3 | 0.171 | 0.226 | 0.216 | 0.232 | 0.280 | 0.285 |
| 0.9 | 10 | 0.264 | 0.312 | 0.331 | 0.360 | 0.284 | 0.409 |
| 0.9 | 30 | 0.313 | 0.356 | 0.356 | 0.418 | 0.317 | 0.441 |
| 0.9 | 100 | 0.331 | 0.363 | 0.362 | 0.440 | 0.343 | 0.480 |
| 0.9 | 1000 | 0.388 | 0.378 | 0.381 | 0.536 | 0.414 | 0.556 |

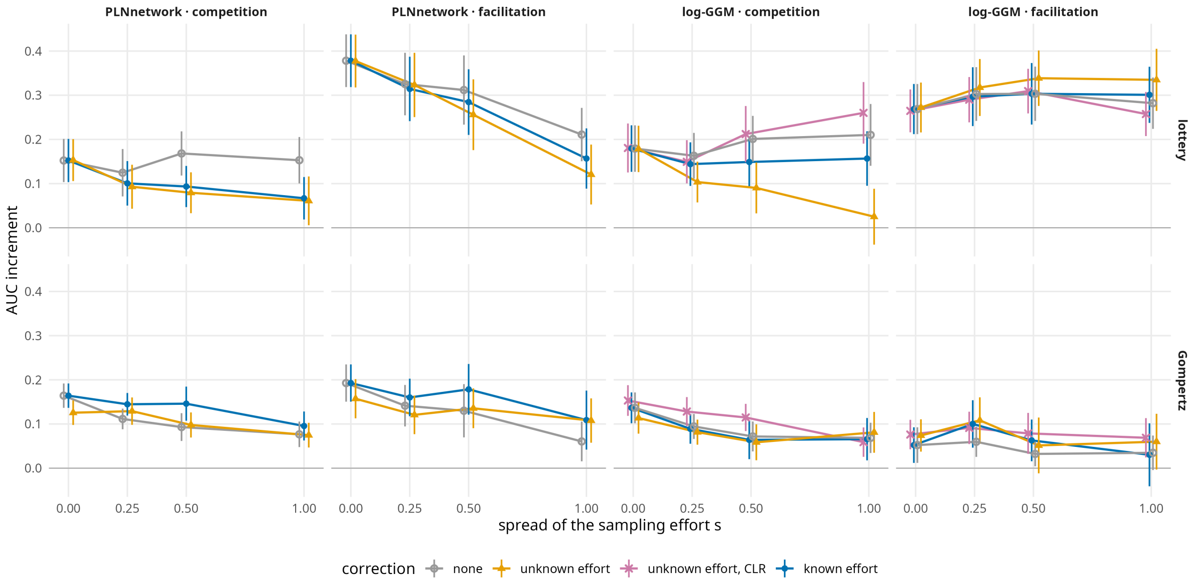


**Figure S12** – Increment of the AUC against the spread $s$ of the sampling effort, at $b = 10$ and $D = 10$; corrections as in Figure 5.

The accuracy of the abiotic estimation does not predict the quality of the network. Under the Gompertz mechanism, the quadratic basis recovers the niche profiles much better than the cubic polynomial and the spline, for the same increment (Table S16). Under the lottery the quadratic basis recovers the niches best and the categorical basis among the worst, yet the second gives the larger increment.

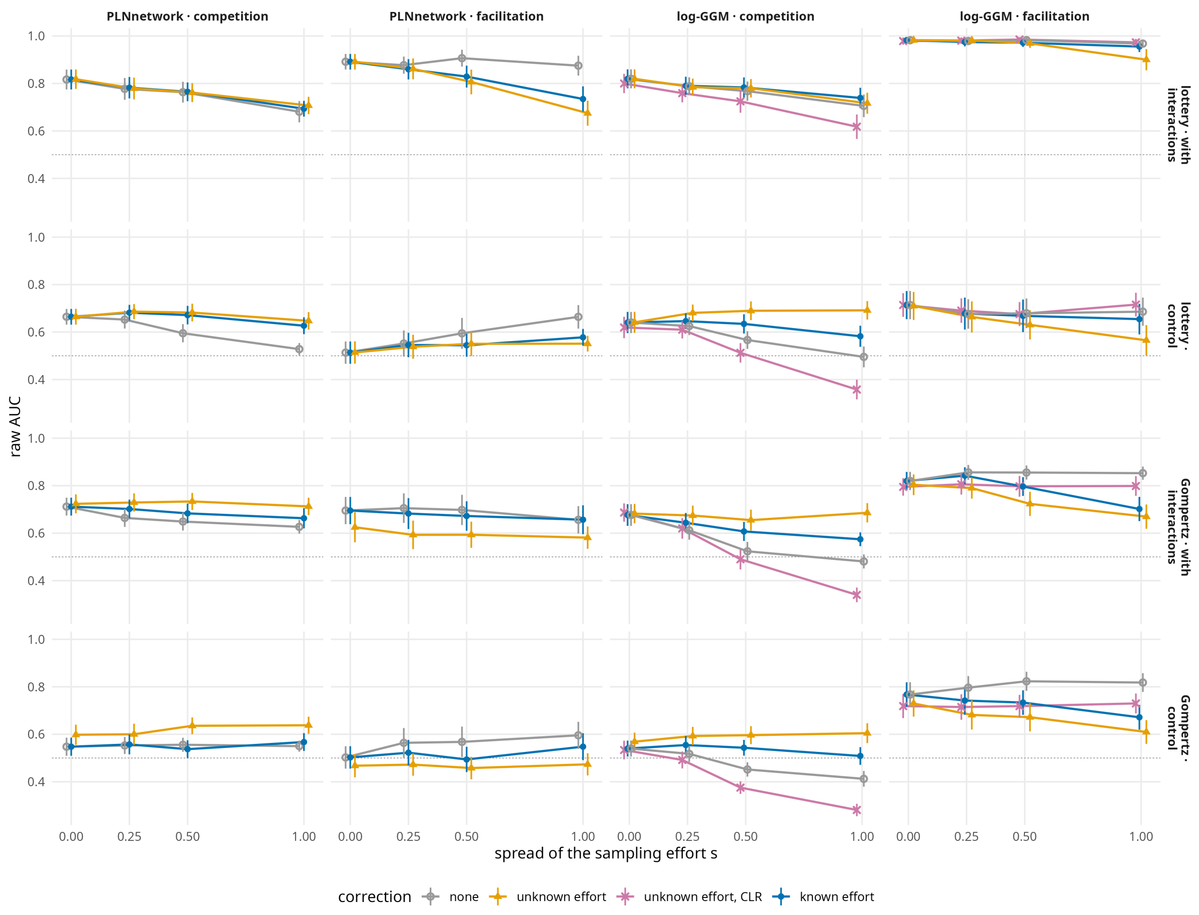


**Figure S13** – Raw AUC of the simulation with interactions and of its paired control against the spread $s$ of the sampling effort, at $b = 10$ and $D = 10$; corrections as in Figure 5. Grey dashes: 0.5, the chance level.

**Table S16** – Quality of the abiotic estimation of the PLN fits, by basis. Niche profile: correlation between the fitted and the true abiotic effect along the gradient, mean over the species. Niche optimum: correlation between the fitted and the true optima. Abiotic share: share of the variance of the latent layer taken by the abiotic effect. Increment: that of PLNnetwork for the AUC of competition with this basis. Means over the settings and the replicates, simulations with interactions.

| mechanism | basis | niche profile | niche optimum | abiotic share | increment |
|---|---|---|---|---|---|
| Lottery | Categorical (10) | 0.41 | 0.49 | 0.89 | 0.12 |
| | Quadratic | 0.92 | 0.98 | 0.93 | 0.10 |
| | Cubic polynomial | 0.62 | 0.98 | 0.94 | 0.09 |
| | Spline (10 df) | 0.33 | 0.98 | 0.97 | 0.06 |
| Gompertz | Categorical (10) | 0.55 | 0.51 | 0.93 | 0.11 |
| | Quadratic | 0.94 | 0.99 | 0.92 | 0.15 |
| | Cubic polynomial | 0.77 | 0.99 | 0.94 | 0.16 |
| | Spline (10 df) | 0.64 | 0.96 | 0.96 | 0.16 |
| Ricker | Categorical (10) | 0.50 | 0.50 | 0.92 | 0.13 |
| | Quadratic | 0.92 | 0.98 | 0.92 | 0.07 |
| | Cubic polynomial | 0.72 | 0.98 | 0.94 | 0.07 |
| | Spline (10 df) | 0.52 | 0.95 | 0.97 | 0.07 |
